# Improving RNA-seq protocols

**DOI:** 10.1101/2025.08.20.671269

**Authors:** Felix Pförtner, Eva Briem, Wolfgang Enard, Daniel Richter

## Abstract

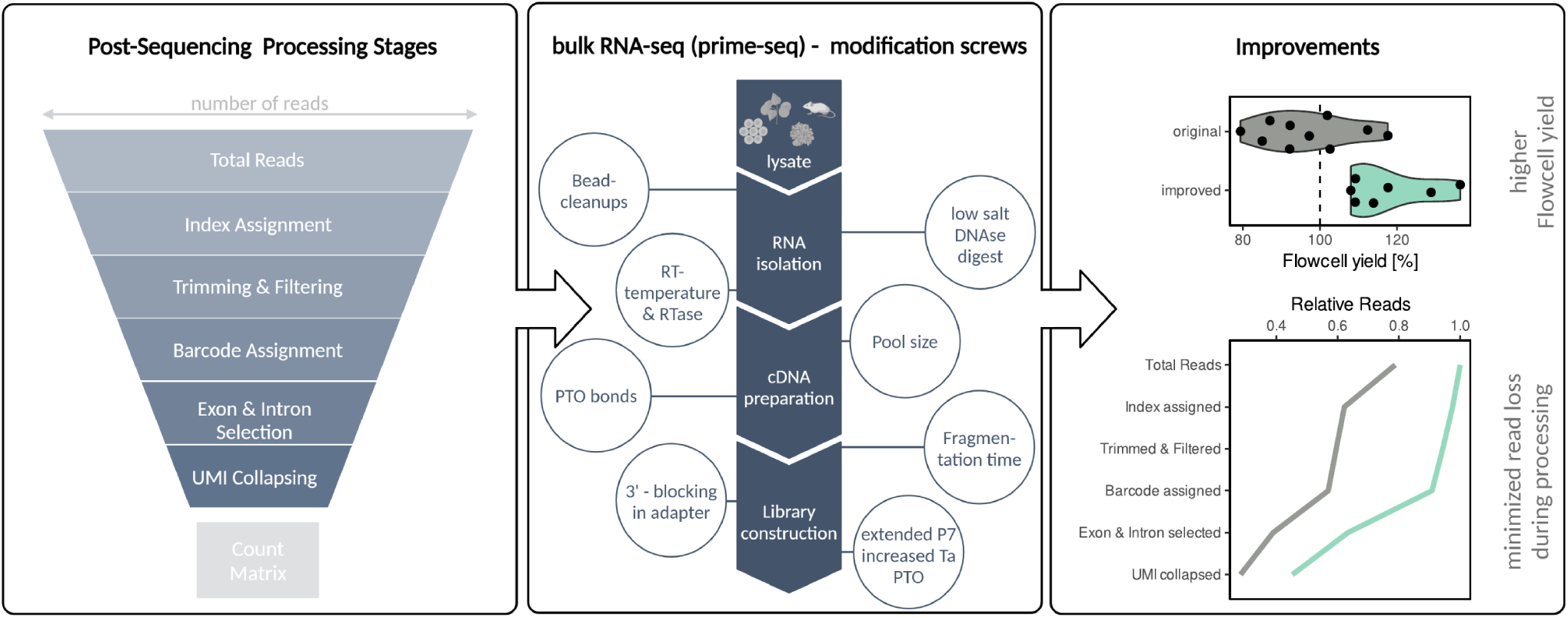

Bulk and single-cell RNA-seq are powerful tools for transcriptomic analysis, providing insights into many aspects of molecular and cellular phenotypes. Costs constrain the amount of biological insight obtainable within a given budget, and as sequencing prices decline, efficient library protocols have become a decisive factor. In this study, we introduce an approach to systematically optimize the number of usable reads that RNA-seq protocols generate. We applied this ”funnel strategy” to prime-seq, an early-barcoding bulk RNA-seq protocol, by systematically testing critical protocol steps totaling 1080 samples in 49 libraries. This resulted in the optimized prime-seq2 protocol that increases the number of usable reads by 60 % and improves one of the most cost-efficient bulk RNA-seq protocols available. Our study also suggests that monitoring the filtering of usable reads can serve as a valuable quality control for many RNA-seq protocols and sheds light on the complexity of the conditions and interactions that shape RNA-seq library composition and their interpretation.

## Introduction

RNA sequencing (RNA-seq) has revolutionized molecular biology by enabling comprehensive analyses of transcriptomes [1]. This includes developments for genome annotation and isoform detection that were boosted by long-read sequencing technologies [2] and comprehensive characterization of cell types, enabled by single-cell RNA-seq and spatial transcriptomics [3]. While these exciting developments have justifiably garnered much attention, bulk RNA-seq, in which average expression levels of many cells are measured, remains an essential and complementary tool. Bulk RNA-seq is valued especially for its cost-efficiency and its easy and unbiased sampling procedure. This is of particular advantage when the cellular composition of samples is sufficiently homogeneous. Notably, the increasing availability of single-cell RNA-seq data, together with recent advances in deconvolution methods [4, 5, 6], is poised to significantly improve the interpretation—and thus the utility—of bulk RNA-seq data in the future.

Many different bulk RNA-seq protocols have been developed [7, 8]. They differ in aspects like the type of RNAs (e.g., polyadenylated RNAs, short RNAs, rRNA-depleted RNAs, newly synthesized RNAs) and the parts of RNAs (5’ ends, 3’ ends, full-length) that get incorporated in sequencing libraries. They differ in their sensitivity, i.e. how many cells are needed as input, their accuracy, i.e. how well cDNA molecules in the library correlate with RNA abundance, their complexity, i.e. how many different RNAs are detected at a given sequencing depth and their technical noise, i.e. the variation with which the detected RNAs are measured in technical replicates. Obviously, different questions and samples require different protocols. Here, we focus on probably the most frequent application of bulk RNA-seq, in which genome-wide expression levels of polyadenylated RNAs are correlated with variables of interest, such as drug treatment, genetic variants, or disease states. Data is usually interpreted by identifying differently expressed genes and their enrichment in molecular pathways and biological processes. This type of bulk RNA-seq application has been called RNA-seq for differential gene expression [1], mRNA Sequencing [9] or quantitative RNA-seq [10]. It is conceptually similar to a quantitative Reverse Transcription PCR (RT-qPCR) for many genes and is becoming as routine, as important, and as affordable as RT-qPCR [11].

As with many methods, costs are a crucial factor, as they limit the amount of biological insight within a given budget. Costs not only constrain the number of replicates and hence the statistical power to detect expression differences [12, 13, 14], but they also limit the number of variables (e.g., tissues, time points, treatments) that can be investigated. For RNA-seq, sequencing costs have long limited the number of samples to the notorious N=3 per condition. Ten years ago, that could be justified by budget limitations when costs for generating and sequencing a bulk RNA-seq sample were maybe around $200. But sequencing costs have dropped ten-fold in the last ten years [15] and one million reads can cost less than one dollar on the latest Illumina NovaSeq X machines. Hence, costs for sequencing an RNA-seq sample fell to $10-20, making costs of $60-164 per sample for commercial library generation kits the dominant cost factor [16]. As a consequence, improvements such as miniaturization and early barcoding, originally developed to improve scRNA-seq cost-efficiency [17], have become increasingly important to leverage the full potential of bulk RNA-seq. Early barcoding protocols (also called Tag-Seq, 3’ RNA-seq, or QuantSeq), in which a sample-specific barcode is introduced either at the 5’ or at the 3’ end of the polyT primed cDNA, allow for pooling cDNA before amplification and library generation. This decreases library reagent costs for non-commercial protocols to $2.53, $4.05, or $6.58 for prime-seq [16], BRB-seq [11], or DECODE-seq [18], respectively. In addition to the reagent costs per library, other factors like necessary equipment, hands-on time, or failure rates contribute to cost-efficiency. Of note, the low costs per library of these protocols make kit-based RNA isolations of ^¼^$5 per sample a relevant factor, and protocols have established different methods to incorporate direct lysis or RNA purification steps [16, 19, 20, 21]. Importantly, costs are directly comparable only at the same efficiency, i.e., at the same statistical power to detect differently expressed genes [22]. At a given number of replicates and sequencing reads, this depends mainly on the library complexity and the technical noise of the protocol [23, 24]. While comprehensive benchmarking for early barcoding protocols has not been done, they generally perform fairly similarly to each other and fairly similarly to commercial gold-standards like TruSeq with respect to technical noise and complexity [16, 11]. A potential drawback of early barcoding protocols is that mainly 5’ or 3’ ends of cDNAs are sequenced, and hence that expression levels are quantified per gene and not per transcript isoform. But as the available functional annotation is almost always gene-specific and not isoform-specific, most projects interpret the RNA-seq data anyway only at the level of differentially expressed genes. Additionally, quantifying expression levels of isoforms requires either long-read sequencing or deep short-read sequencing, which increases the sequencing costs per library considerably. Hence, the most cost-efficient way to quantify gene expression levels by RNA-seq is currently early barcoding RNA-seq.

So while the cost-efficiency of bulk RNA-seq has made tremendous progress by reducing reagent costs per library, relatively little attention has been devoted to quantifying and optimizing how many usable reads, i.e., reads that are used to quantify gene expression levels, are actually produced per library. In general, RNA-seq libraries can differ enormously in that respect, as e.g., seen by an analysis of *>*2000 tumor bulk RNA-seq datasets across *>*40 studies in which the fraction of usable reads ranged from 0 % to 79 % at a median of 50 % [25]. Early barcoding protocols show similar patterns with substantial variation among protocols and among samples. For instance, increasing the number of usable reads was a main motivation to develop BRB-seq from the previous SCRB-seq protocol [11] , and when using prime-seq (another SCRB-seq derivative), we also observed substantial variations in the number of usable reads among samples and especially among sample types. In particular, we observed that some sequencing runs on patterned flow cells of Illumina’s NextSeq generated fewer usable reads and especially fewer total reads than expected. We had not observed this before on non-patterned flow cells such as Illumina’s Hi-Seq, probably because on those flow cells, one can ignore clustering molecules that do not generate sequencing reads when overloading flow cells.

To optimize the number of usable reads for prime-seq we systematically varied different steps in the protocol. In this way, we could increase the number of usable reads by 60 %, importantly, without compromising the complexity and noise of the prime-seq libraries. This new version - prime-seq2 - is a considerable improvement in cost-efficiency, reducing sequencing costs per sample, e.g., from $53.1 to $33.2, when generating 10 million usable UMIs at $1.50 per million reads, at an unchanged cost of ∼$3.5 per library including RNA isolation. As we think that many commercial or home-brew early barcoding protocols could profit from such an optimization, we present our approach as a generally applicable optimization strategy that visualizes the amount of usable reads at different processing steps as a funnel (Fig. 1A).

**Figure 1.**
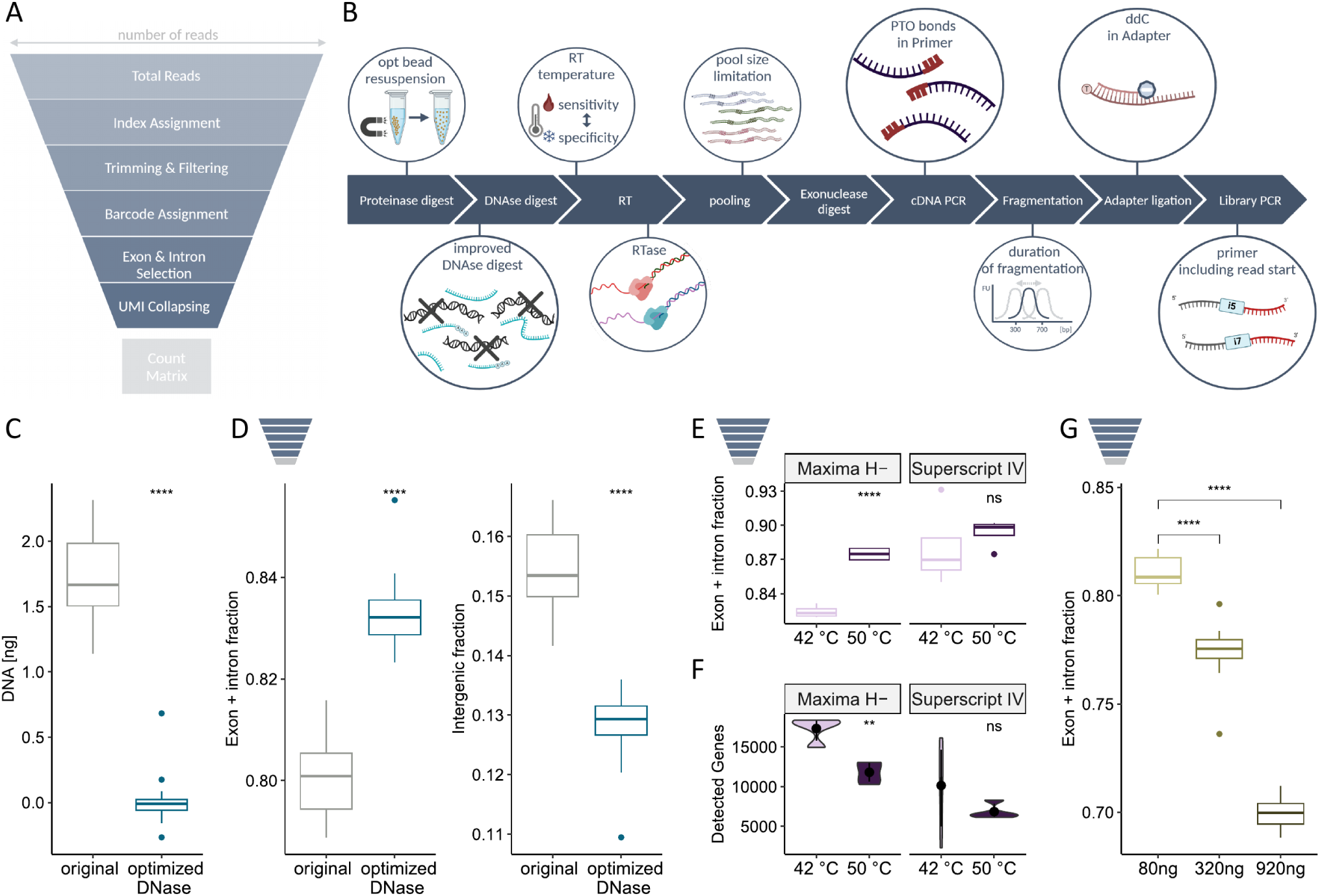
The funnel strategy and its application to prime-seq. **A**, Funnel strategy scheme. The total reads from the sequencer - having per definition at least a Read1 sequence - are filtered for the presence of both Illumina indices, for the presence of long high quality reads, for the presence of a barcode introduced with the RT-primers, for reads mapping to the coding strand of annotated exons and introns and for reads that differ in their UMI sequence and hence are not PCR duplicates. These UMI counts per gene and per sample constitute the count matrix that is used for all subsequent analysis and hence is the number of usable reads. **B**, Overview of the prime-seq workflow and the parameters that have been tested in this study. For a more detailed graphical version, see Figure S7. **C**, DNA amount measured after the original DNase treatment under high salt conditions on beads and under the optimized conditions in solution (n=64; Two independent replicates of 16 samples each per condition.) **D**, The fraction of reads mapping to exons and introns (left) and intergenic regions (right) indicate that the optimized DNA digest removes remaining genomic DNA (n=32; 16 samples per condition in one library.) **E,** The fraction of reads mapping to exons and introns is lower in the original RT condition (Maxima H- at 42 °C) than in the other tested conditions (lines are medians of n=4 per condition). **F,** The number of detected genes, downsampled to 7 million reads per sample is higher in the original RT condition (Maxima H- at 42 °C) than in the other tested conditions (dots are averages of n=4 per condition). **G**, The fraction of reads mapping to exons and introns decreases strongly with increasing amounts of samples pooled after reverse transcription. Samples à 10 ng of total RNA were reverse transcribed, pooled, exonuclease digested, and then added to the cDNA amplification PCR in amounts corresponding to 8 (80 ng), 32 (320 ng), and 92 (920 ng) samples (total RNA), respectively. This was done in a total of 40 replicates (n=8 for 80 ng and n=16 for 320 ng and 920 ng) from which one library per condition was generated P-values were calculated in unpaired two-sided t-tests (ns: p *>* 0.05; *: p ≤ 0.05; **: p ≤ 0.01; ***: p ≤ 0.001; ****: p ≤ 0.0001).

## Materials and Methods

### Preparation of SPRI Beads

Homemade solid-phase reversible immobilization (SPRI) beads are used throughout the paper. They are produced as follows: the 22 % PEG solution (22 % (w/v) PEG 8000, 1 M NaCl, 10 mM Tris-HCl pH 8, 1 mM EDTA. 0.01 % IGEPAL, 0.05 % sodium azide, to 49 ml with UltraPure (UP) H_2_O) was heated at 40 °C to dissolve PEG. Sera-Mag Speed Beads (1 ml) were separated from their solution using magnets. The supernatant was removed. The resulting pellet was washed twice in TE buffer (10 mM Tris-HCl, pH 8, 1 mM EDTA) and eluted in 0.9 ml TE buffer. This bead suspension was mixed with the 22 % PEG solution. SPRI beads were stored at 4 °C and equilibrated to room temperature before usage.

### Nucleic Acid Purification using SPRI Beads

Home-made SPRI beads were used for nucleic acid purification at multiple steps during the RNA-seq protocol. In the ensuing procedures, ”cleanup” always followed this protocol: Bead suspension was added in the specified ratio, incubated for 5 min, and the beads were anchored using magnets. The supernatant was discarded, and the beads washed twice with 50-1000 *µ*l 80 % ethanol in UP H_2_O. Beads were air-dried, removed from the magnet and eluted in UP H_2_O. After 5 min incubation, beads were anchored again using magnets and the supernatant was transferred to a new well.

### Sample Preparation

To compare different RNA-seq protocols on the same source material, an abundant and representative tissue or cell lysate was needed. Brain tissue from a 250-day-old male *Foxp2 ^hum/hum^* mouse (5H11 line [26]) was used. The tissue was homogenized in RLT Plus with 1 % *β* -mercaptoethanol and stored at -70 °C. For validation, a second, more challenging sample was used, which delivered worse results with the original protocol. Striatal punches of C57BL/6J mice were lysed in RLT Plus with 1 % *β* -mercaptoethanol and stored at -70 °C. Total RNA concentrations were determined using the QuantiFluor RNA System after Proteinase K treatment, DNA digest and purifications using SPRI beads.

### Standard Prime-seq Workflow

The RNA-seq workflow used as a starting point is prime-seq [16]. It is termed as the “original” prime-seq method and used in this way if not mentioned otherwise. It is described in the following section and visualized in Fig. S7. The full list of reagents and oligonucleotides can be found in Tab. 1 & 2.

**Table 1.**
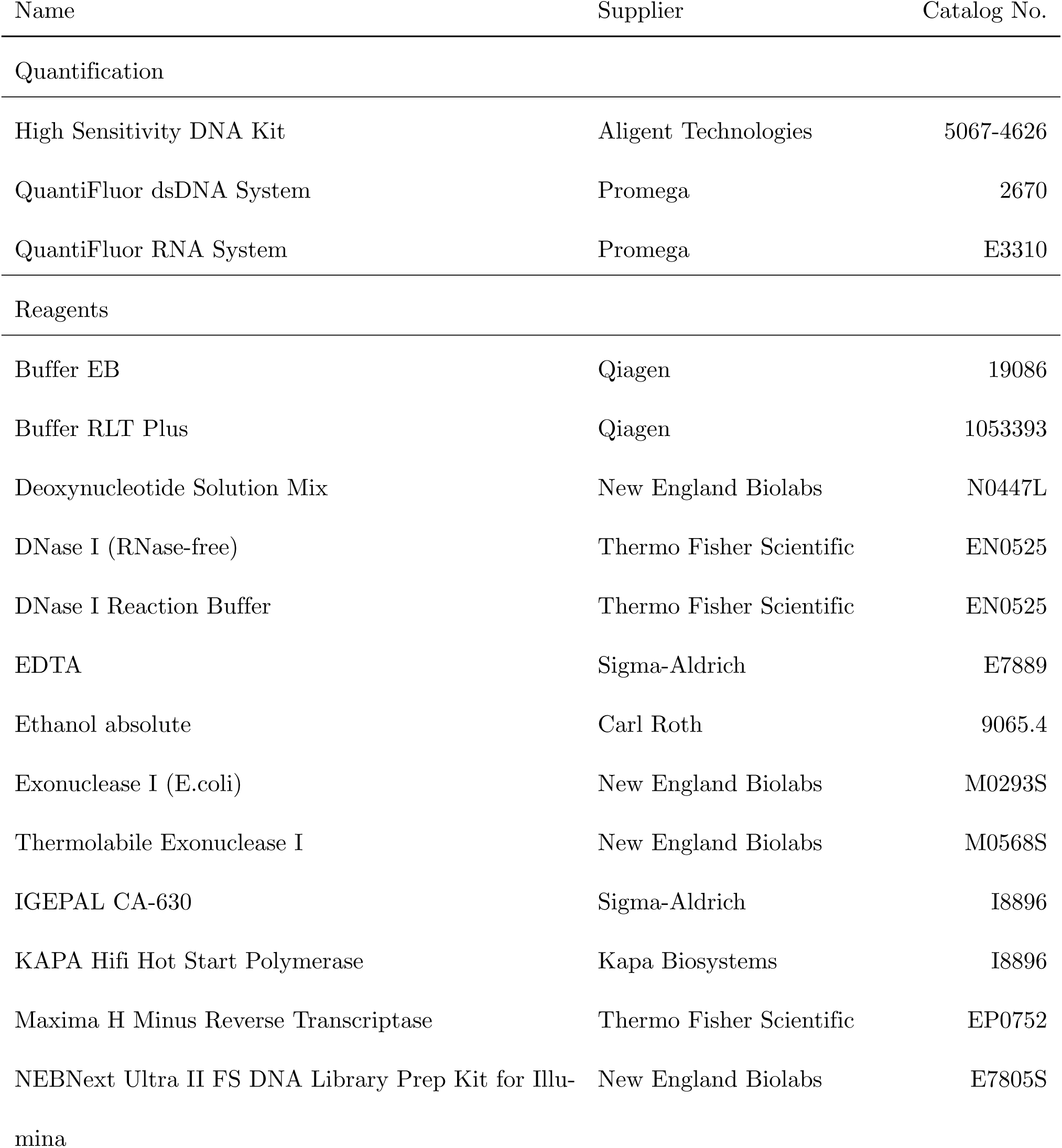
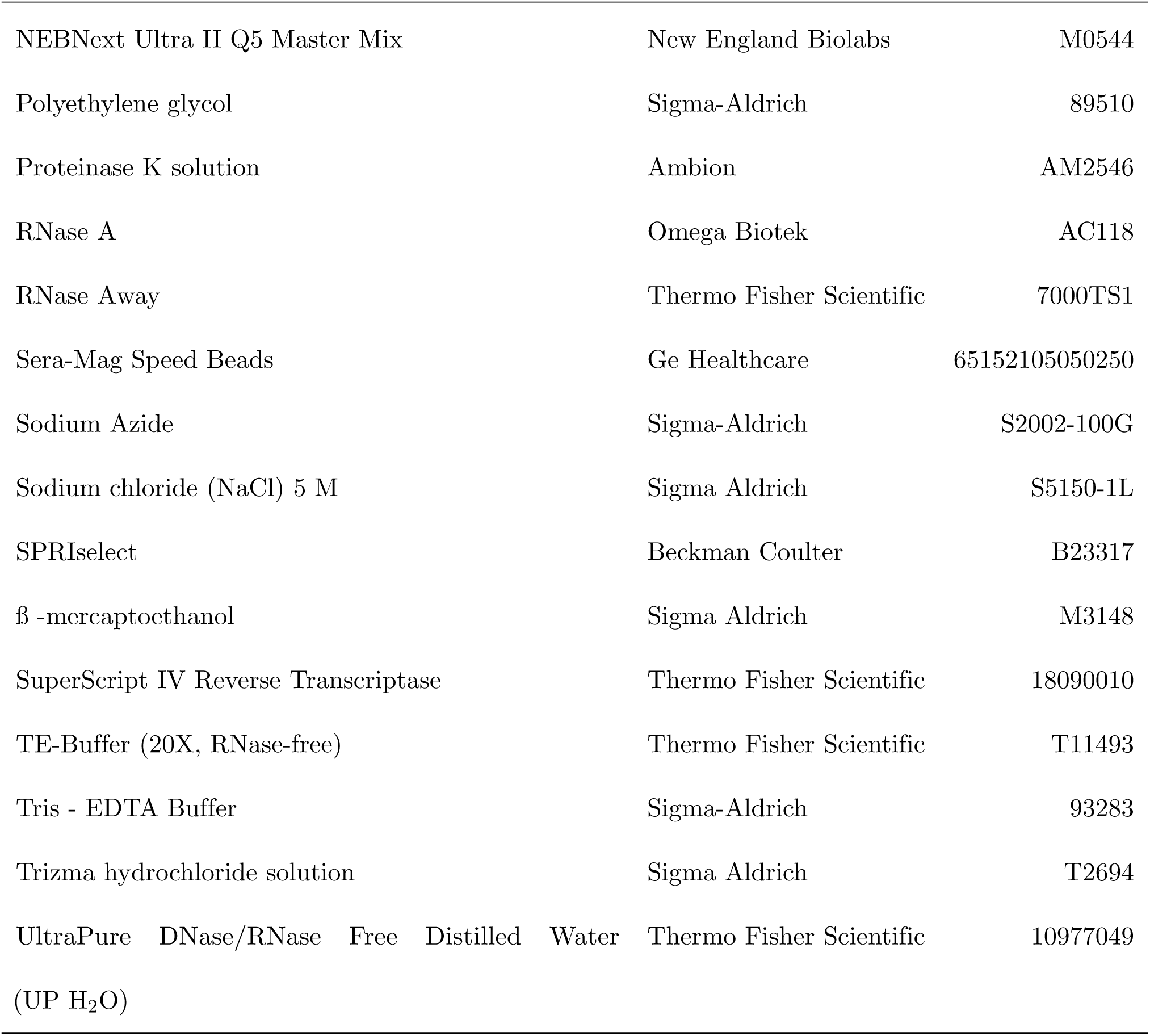
Enzymes & Chemicals.

**Table 2.**
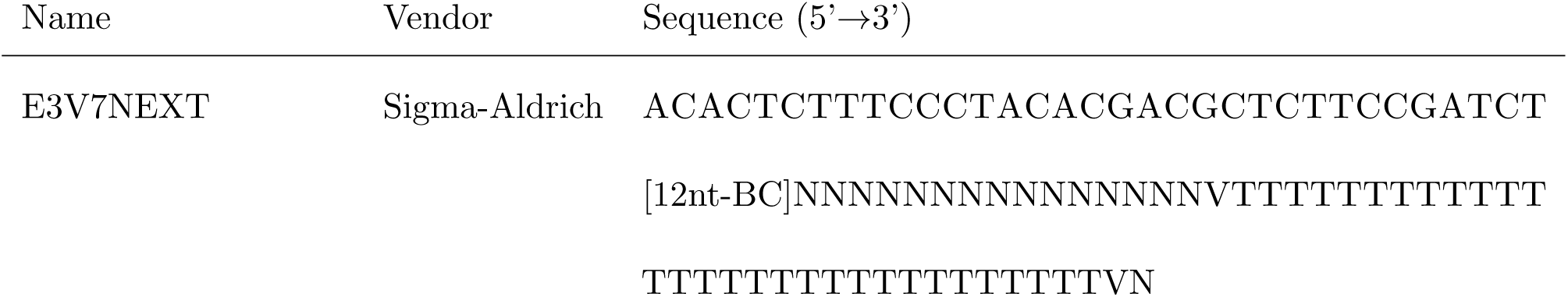
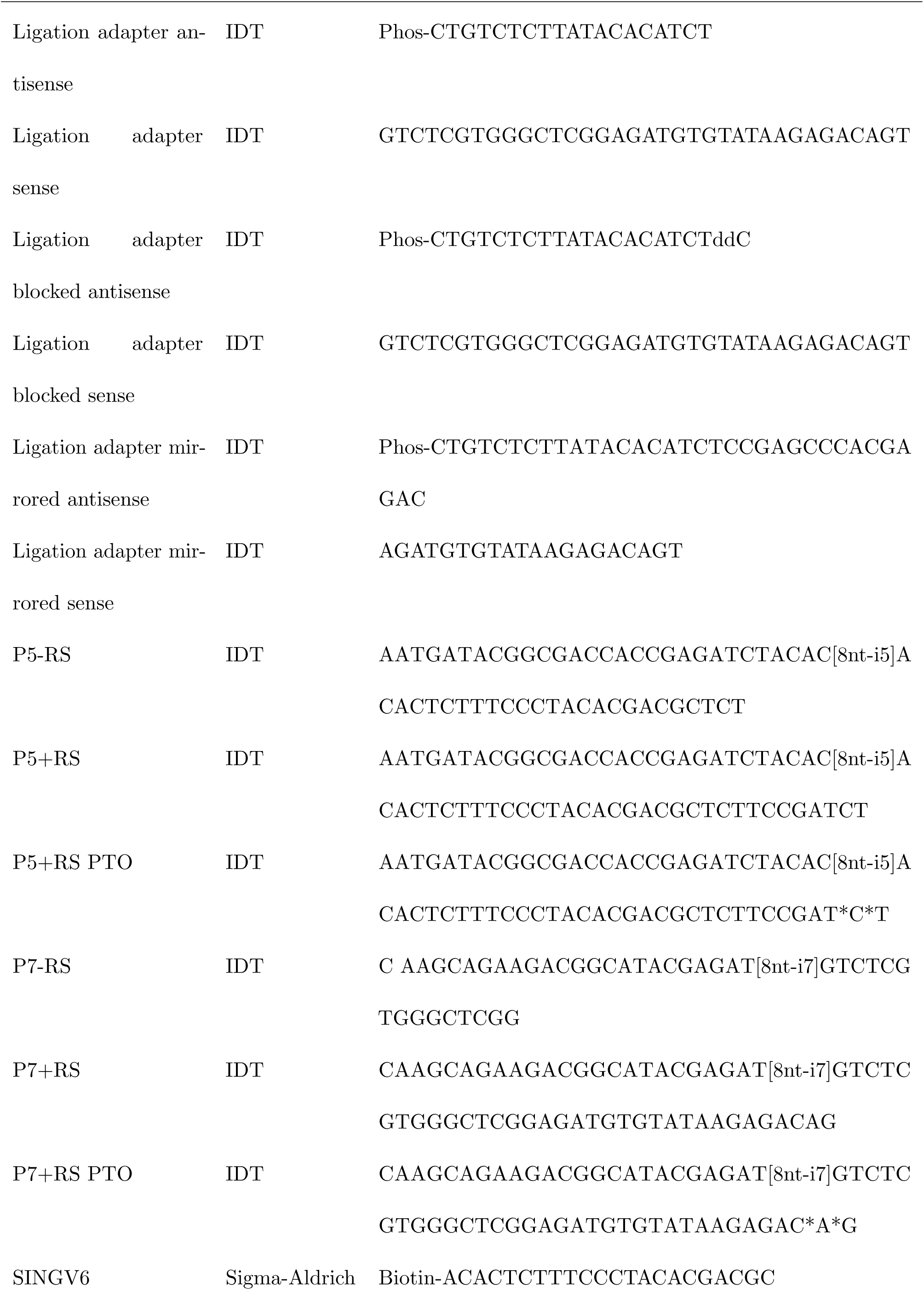
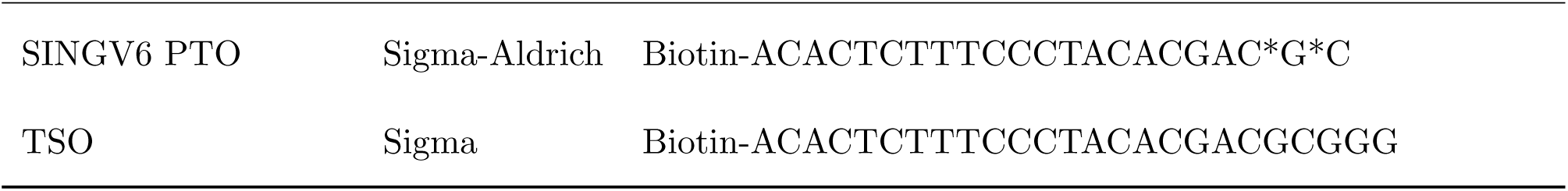
Oligonucleotides used in this study. ”Phos” signifies a 5’ phosphorylation. ”*” signifies a PTO bond. E3V7NEXT contains a region (UMI) and an anchor of random bases denoted by ”N”. Regions in square brackets are barcodes (BC) or indices (i5 & i7) with varying sequences.

#### cDNA Preparation

Once the RNA concentration of the standard lysate sample had been established, the standard input of the samples was adjusted to target approximately 10 ng of total RNA per sample. First, proteins were digested using 20 *µ*g Proteinase K in 0.5 mM EDTA at 50 °C for 15 min, followed by heat inactivation at 75 °C for 10 min. SPRI beads were used in a 2× ratio (given as bead volume to sample volume) to purify the samples as described above.

In this step, samples were resuspended in 5 *µ*l UP H_2_O without removing the beads. DNA was digested on beads using 1 unit of DNase I at 20 °C for 10 min in 1× DNase I buffer and 1× bead binding buffer (11 % (w/v) PEG 8000, 0.5 M NaCl, 5 mM Tris-HCl pH 8, 0.005 % IGEPAL, 0.025 % sodium azide). EDTA was added to 4.5 mM before heat-inactivation at 65 °C for 5 min. The samples were cleaned up again and eluted in 4 *µ*l UP H_2_O. RT was performed in 1× Maxima H Minus buffer using 30 units Maxima H Minus, 1 mM each dNTP, 1 *µ*M template-switching oligo (TSO), and 1 *µ*M barcoded oligo (dT) primers (E3V7NEXT). The RT reaction was incubated at 42 °C for 90 min. The samples were combined, purified using SPRI beads at a 1× ratio and eluted in 17 *µ*l in UP H_2_O. This “pooled” sample was termed the “preAmp pool”. Residual primers were digested using 20 units Exonuclease I in 2 *µ*l 10× Exonuclease I buffer at 37 °C for 20 min with ensuing heat inactivation at 80 °C for 10 min.

The samples were cleaned using SPRI beads at a 0.8× ratio, and eluted in 20 *µ*l of UP H_2_O. cDNA was amplified in 1× KAPA HiFi Ready Mix and 0.6 *µ*M SINGV6 primer, according to this PCR protocol: initial denaturation at 98 °C for 3 min, denaturation at 98 °C for 15 s, annealing at 65 °C for 30 s, elongation at 68 °C for 4 min, and final elongation at 72 °C for 10 min. Denaturation, annealing, and elongation were repeated for 10–23 cycles, depending on the initial amount of total RNA. The DNA was cleaned with SPRI beads in a 0.8× ratio and eluted with 10 *µ*l of UP H_2_O. The cDNA concentration was measured using the QuantiFluor dsDNA System and the quality of the cDNA was assessed using a Bioanalyzer.

#### Library Preparation

Libraries were prepared using the NEBNext Ultra II FS Library Preparation Kit. 1.2-4 *µ*l cDNA (usually 4–20 ng) were fragmented using the FS Enzyme Mix and FS Reaction Buffer in a 6 *µ*l reaction by incubating at 37 °C for 5 min, then at 65 °C for 30 min. 0.5 *µ*l ligation adapter (1.5 *µ*M) was ligated using the Ligation Master Mix and Ligation Enhancer in a reaction volume of 12.7 *µ*l by incubating at 20 °C for 15 min. Double-sided size selection was performed using SPRIselect beads according to the manufacturer, with a bead ratio of 0.52× for the removal of long DNA fragments and a ratio of 0.72× for the removal of short fragments. Samples were eluted in 11 *µ*l 0.1× TE buffer. 10.5 *µ*l DNA sample was amplified using 12.5 *µ*l Q5 Master Mix, 1 *µ*L Nextera P7 index primer (5 *µ*M), and 1 *µ*L TruSeq P5 index primer (5 *µ*M) following this PCR protocol: initial denaturation at 98 °C for 30 s, denaturation at 98 °C for 10 s, annealing and elongation at 65 °C for 1 min 15 s, and final elongation at 65 °C for 5 min. Denaturation and annealing/elongation were repeated for 10–13 cycles, depending on the initial amount of cDNA. Double-sided size selection was performed as above. The initial DNA concentration was measured using the QuantiFluor dsDNA System and, more precisely, using a Bioanalyzer, which also allowed for quality assessment.

### Prime-seq Interventions

The following experiments are all modifications of the original prime-seq protocol described above.

#### Cleanup Resuspension

The cleanup steps following Proteinase K and DNase I digest were altered to include resuspension in ethanol. After adding 80 % ethanol, the plate was taken from the magnet, beads were resuspended in ethanol by pipetting, beads were reanchored using the magnet and ethanol was removed. This procedure was repeated in the second wash step.

#### DNase I Digest

In order to decrease the NaCl concentration during DNase I digest, bead binding buffer was removed from the reaction, and UP H_2_O was added to keep the reaction volume constant. Consequently, 18 *µ*l bead binding buffer was added after heat-inactivation to obtain the same PEG concentration as with the initially added bead binding buffer.

#### Reverse Transcription

The incubation temperature during reverse transcription (RT) was increased from 42 °C to 50 °C. The number of PCR cycles during preAmp PCR was increased by six cycles at 50 °C to obtain similar cDNA yields. Additionally, Superscript IV was tested against the standard Maxima H Minus as an alternative reverse transcriptase (RTase). In this reaction setup, Maxima buffer was swapped for Superscript IV buffer, and DTT was added to 5 mM, while keeping all other reagents and the total volume constant.

#### cDNA amplification pool size

To test the effect of combined RNA input in the cDNA amplification, pools of 8, 32, and 92 samples were compared. Sequencing data was downsampled to equal read numbers for analysis. For length binning, each gene was represented by a single transcript selected in the following hierarchical order: (1) the consensus coding sequence (CCDS), (2) the transcript with the highest support level, and (3) the longest transcript. To determine anti-sense intergenic reads, all intergenic reads were recounted using zUMIs (v2.9.7 [27]) with library strandedness set to reverse.

#### Fragmentation

To determine the optimal fragmentation time, cDNA from standard brain lysate was processed in the standard prime-seq library preparation without size selection. 5, 15, and 80 ng cDNA were fragmented for 0, 2, 3.5, 4, 4.5, 5, 6, 7, 8, or 15 min. Fragmentation was compared using a Bioanalyzer.

#### Ligation Adapters and Index primers

Altered ligation adapters and index primers were compared. The mirrored ligation adapter contains swapped strands to have a 3’ overhang instead of a 5’ overhang. The blocked prime-seq ligation contains an added dideoxycytidine at its 3’ end. The P5+RS and P7-RS primers correspond to the primers used in the original prime-seq protocol. The P5-RS primer was shortened by 8 nt from the 3’ end, which decreases its melting temperature from 75 °C to 71 °C. The P7+RS primer was extended by 19 nt at the 3’ end to its maximum length, which increases its melting temperature from 65 °C to 74 °C. Only the P5/7+RS primers contain the complete start sites for priming of the read 1 and read 2 sequencing reads, respectively. In combinations of P5+RS and P7+RS, Annealing/Elongation temperature during library PCR was increased from 65 °C to 72 °C, as calculated using the NEB Tm calculator.

#### PTO bonds

In the cDNA PCR and the library PCR, SINGV6 and index primers (P5+RS and P7+RS) with two 3’-terminal phosphorothioate (PTO) bonds were used.

### Sequencing

RNA-seq libraries were sequenced by the Blum Lab as part of the Laboratory for Functional Genome Analysis (LAFUGA) of the Gene Center Munich. Sequencing was performed on an Illumina NextSeq 1000/2000 instrument with the following setup: Read 1: 28 bp, Index 1: 8 bp, Index 2: 8 bp, Read 2: 93 bp.

### DNase digest test

DNase digest was compared using the original protocol, the improved protocol, and a negative control lacking DNase I. 8 samples per condition of mouse liver lysate containing 50 ng total RNA each were used. After Proteinase K treatment, the DNase I digest was done as in the original protocol, without DNase I and using the improved low-salt conditions. RNA was quantified using the QuantiFluor RNA System. RNA was digested using 1 *µ*l RNase A per sample at room temperature for 90 min. DNA was quantified using the QuantiFluor dsDNA system after a bead clean-up. The experiment was independently repeated.

### Data Analysis

Prime-seq libraries were demultiplexed with deML (v1.1.1 [28]) using the indices in the index primers. Terminal poly(A)s were trimmed from read 2 and read pairs were filtered to minimum lengths of 28 nt read 1 and 40 nt read 2 using cutadapt (v3.2 [29]): ”cutadapt -A A{30} –discard-casava –minimum- length 28:40”. zUMIs (v2.9.7 [27]) was used for further processing. Reads were filtered if more than 2 barcode bases had Q-scores below 20 or if more than 3 UMI bases had Q-scores below 20. Barcodes (read 1, 1-12 nt) were demultiplexed and binned within a Hamming distance of 2. Remaining reads were mapped to the genome GRCm39. Exonic and intronic reads and UMIs (read 1, 13-28 nt) were counted per gene according to Gencode version 34.

Power simulations were performed with powsimR (v1.2.5 [30]). For each protocol and sample size setup (3 vs. 3, 6 vs. 6, 12-20 vs. 12-20, 24 vs. 24, and 48 vs. 48), 25 simulations were performed with a negative binomial distribution, trimmed mean of M values (TMM) normalization, limma-voom DE-method, and 10 % differentially expressed genes. The log fold change was modeled by a gamma distribution with shape = 1 and rate = 2. The data were stratified by absolute log fold change or mean expression.

Further analysis was done in R (v4.2.3 [31]) using tidyverse (v2.0.0 [32]), data.table (v1.14.8 [33]), ggbeeswarm (v0.7.2 [34]), purrr (v1.0.2 [35]), ggpubr (v0.6.0[36]), patchwork (v1.1.3 [37]), cowplot (v1.1.1[38]), ShortRead (v1.56.1 [39]) and bioanalyzeR (v0.10.1 [40]).

### Reagents

## Results

### A funnel strategy to systematically optimize the number of usable reads for bulk RNA-seq protocols

When sequencing RNA-seq libraries, a certain number of reads per sample are targeted, often between 5 and 20 million reads, depending on the number and types of replicates and the intended analyses. After pooling and quantification, one loads the required amount of library molecules on the flow cells of short-read sequencers such as Illumina’s NextSeq or NovaSeq. This sets the price tag, but how many reads one gets from the flow cell and how many of those can be used for quantifying expression levels depends largely on the molecules in the library and, hence, largely on the RNA-seq protocol. For example, reads that cluster on the flow cell but lack a functional read primer binding site will lead to fewer reads than expected. Similarly, molecules lacking indices or barcodes, having inserts not derived from mRNAs or PCR duplicates (identified by having the same Unique Molecular identifier - UMI), will be filtered out and reduce the number of usable reads that go into the count matrix, i.e. the table in which reads are counted per sample (columns) and gene (rows). Note that we use reads mapping to exons and introns of genes, as it has been shown that intronic reads are generally derived from pre-mRNA and help to quantify gene expression levels [41, 16, 27]. Our goal was to increase the cost-efficiency of our prime-seq protocol by increasing the number of usable reads without compromising the complexity of the libraries. To visualize bottlenecks, we found it helpful to use a ”read funnel” displaying the number of usable reads retained after each sequential filtering step (Fig. 1A), a strategy allowing a better visualization and systematic optimization of RNA-seq protocols in general.

### Identifying potential optimizations for prime-seq

We had previously developed and benchmarked prime-seq and could show that its power is comparable to TruSeq, making it fourfold more cost-efficient due to almost 50-fold cheaper library costs [16]. It uses the principles of poly(A) priming, template switching, early barcoding, and UMIs to generate 3’ tagged RNA-seq libraries (Fig. 1B and Fig. S7) and hence is similar to 3’ early barcoding bulk RNA-seq protocols like BRB-seq [11] or DRUG-seq [20] and to the 3’ early barcoding scRNA-seq protocol of 10x Genomics.

Prime-seq has been used extensively in our lab over the past years. During that time, and especially with the switch from non-patterned flow cells of Illumina’s HiSeq to the patterned flow cells of the NextSeq, we noticed that, for some samples, sequencing yields were 20–30 % lower than expected based on Illumina’s nominal flow cell output specifications. We also noticed that we often lost a considerable (10-30 %) amount of reads for which we couldn’t assign the i7 index, and that some sample types had a very high proportion of intergenic reads (*>*30 %). These patterns were not immediately apparent, as they had always been present to some degree and varied substantially across samples. Furthermore, it is not obvious how these patterns are caused and how they could be fixed, given the complexity of interactions among the different protocol steps and sample types.

As even small improvements could provide considerable benefits for a highly used protocol and as testing variations might reveal general mechanistic insights into the generation of complex sequencing libraries, we tested nine protocol steps that we identified as promising for increasing the number of usable reads (Fig. 1B). We iteratively tested these protocol modifications, amounting to a total of 1080 RNA-seq samples in 49 libraries for a total of 131 million reads that we structured and visualized using our funnel strategy. For initial testing, a mouse whole-brain lysate was used as an abundant, homogeneous sample. Later, lysates from mouse striatal punch biopsies were used as more challenging samples for which we had previously seen a very low fraction of usable reads with the old protocol. In the following, we discuss the tested modifications in the order they appear within the protocol.

### Additional RNA washing increases exonic reads but lowers complexity

RNA extraction from lysates comprises the digestion of proteins using Proteinase K, followed by a DNase treatment. After each step, samples are purified using SPRI beads to remove substances such as salts, small nucleic acids, proteins, lipids, or polysaccharides that can negatively impact subsequent steps in the protocol. This is typically done by adding ethanol for a few seconds to the immobilized beads on the magnet. We reasoned that a resuspension of the beads in ethanol at the washing steps might enhance the purification process by removing more contaminants and, in this way, increase the number of usable reads. This additional resuspension increased the mean proportion of exonic reads from 68 % to 74 % while reducing intronic, intergenic and unmapped reads (Fig. S1A). However, additional resuspension also led to a reduction in the mean number of UMIs per read by 2 % (Fig. S1B). We also observed a non-significant 7 % decrease in the library complexity, i.e. in the number of detected genes at 1.25 million reads per sample with a larger spread of detected genes among samples (Fig. S1C).

Hence, the additional RNA washing increased exonic reads, but led to fewer usable reads in the end and to less complex and potentially more variable libraries. We do not know the reasons for these changes, but as the additional washing does not improve the number of usable reads and as it increases hands-on time, we decided not to incorporate this additional washing step into an improved prime-seq protocol. However, it might remain an option worth testing for lysates with more challenging compositions than whole brain lysates, such as samples with a very high lipid content or chlorophyll-containing samples.

### Improved DNase treatment reduces intergenic reads

Funnels from previous prime-seq projects indicated that a considerable fraction of reads is lost at exon and intron assignment due to 10-20 % of reads mapping to intergenic regions within the genome, i.e. all reads not mapping to the expected strand of the current version of GENCODE annotated exons and introns [27]. Reverse transcriptases used for cDNA generation, including the engineered MMLV-type RTs such as Maxima and Superscript, can generally also use single-stranded DNA as a template [42] and hence these reads can in principle be derived from contaminating genomic DNA. However, our previous experiments with mixing human total RNA and mouse genomic DNA had shown that the fraction of UMIs derived from mouse genomic DNA is below 5 %, even if more than half of the input is mouse DNA and no DNase I treatment is done [16]. Furthermore, the fraction of UMIs derived from mouse genomic DNA is ∼ 0 %, even if more than half of the input is mouse DNA and a DNA digest is done, although this digest is done under suboptimal high salt conditions on beads [16]. These findings are difficult to reconcile with a high fraction of intergenic reads derived from genomic DNA. However, one potential explanation could be the different behavior of an intact, high molecular weight DNA present in the mouse-human test samples and a more degraded, low molecular weight DNA in problematic samples that show a high number of intergenic reads. Larger DNA fragments would be expected to stick more firmly to the beads and, consequently, could be less accessible for reverse transcription than smaller fragments. Consequently, samples with fragmented genomic DNA in combination with a partial DNA digest due to suboptimal conditions on the beads could lead to more accessible DNA and hence to more intergenic reads. In contrast, under optimal conditions, DNase I should be able to digest the entire genomic DNA to di-, tri-, and short oligonucleotides [43], which are then removed during bead clean-up. Hence, we tested whether an optimized DNA digest would lead to less DNA and to fewer intergenic reads. For this, we compared conditions without DNase I treatment, with the original DNase I treatment under suboptimal (high-salt) conditions on beads, and with the optimized DNase I treatment under low-salt conditions off beads. Indeed, when performing the prime-seq RNA isolation protocol on the mouse whole brain samples, the DNA content in the purified lysate was reduced from 1.7 ± 0.4 ng DNA (mean ± SD) for the original DNase I treatment to 0.0 ± 0.2 ng DNA for the optimized treatment (Fig. 1C). This indicates strongly that the original protocol indeed left residual DNA after DNA digestion, which was eliminated in the optimized digest that lowered RNA content only minimally (Fig. S2A). Interestingly, samples without any DNase I treatment contained 0.3 ± 0.2 ng of DNA, i.e., considerably less than for the original DNase I treatment. This supports that high molecular weight DNA indeed binds strongly to beads and hence cannot be eluted very well.

To assess the impact of the improved DNase conditions on the final data, we produced libraries from the mouse brain lysates with and without this optimization. As hypothesized, this reduced the fraction of intergenic reads compared to the original protocol from 16 % to 13 % and accordingly increased the fraction of exonic and intronic reads from 80 % to 83 % (Fig. 1D). Assuming that the optimized DNA digest is complete, as supported by the absence of DNA after elution and as expected from the DNase I activity under these conditions, the remaining intergenic reads must be derived from reverse transcribed RNA (see discussion below).

In summary, we were able to optimize the genomic DNA digestion within the protocol without additional costs. For our mouse brain samples, this increased the number of usable reads by 3 %, but we expect that the improvement can also be considerably bigger for samples that contain more DNA and more partially degraded DNA. Hence, we integrated this optimized DNA digest into our protocol.

### Reverse transcription as a trade-off between complexity and specificity

Reverse transcription (RT) is the crucial step in any RNA-seq workflow. To prime reverse transcription of mRNAs, prime-seq currently uses an oligo(dT)_30_ primer with a VN-anchor and the RTase Maxima H- at 42 °C. However, the manufacturer suggests a reaction temperature of 50 °C for this enzyme when utilizing oligo(dT)_18_ primers. Increasing the temperature during reverse transcription is expected to enhance priming specificity and to facilitate the denaturation of RNA secondary structures, thereby improving access to more mRNAs [44]. Additionally, other RTases could improve specificity and complexity of the generated cDNAs, such as the generally high-performing Superscript IV [45, 46, 47, 48]. To test the influence of the RTase as well as the RT temperature, we tested both RTases, Maxima H- and Superscript IV, at 42 °C and 50 °C with the mouse brain lysate. This resulted in 6 % and 4 % more exonic and intronic reads when using Maxima H- and Superscript IV, respectively. Similarly, substituting Maxima H- with Superscript IV resulted in 4 % and 2 % more exonic and intronic reads at 42 °C and 50 °C, respectively. In total, switching from Maxima H- at 42 °C to Superscript IV at 50 °C elevated the mean exonic and intronic read fraction from 82 % to 89 % (Fig. 1E).

However, these enhancements also led to a decreased number of detected genes. At a sequencing depth of 3 million reads per sample, increasing the temperature alone reduced the mean number of detected genes by 32 %. Switching the enzyme to Superscript IV further decreased the mean number of detected genes by 42 % (Fig. 1F).

So while we see an improvement in the number of usable reads, this increase in specificity comes at the cost of library complexity. This trade-off illuminates the importance of considering not only one quality metric when optimizing a protocol. As we think that a gain of 8 % in usable reads when increasing temperature and switching RTase is not worth a loss of 60 % of detected genes, we decided to stick with using Maxima H- at 42 °C.

### Pooling fewer samples increases the number of usable reads

The key advantage of early barcoding methods is the pooling of cDNA after Reverse Transcription for PCR amplification and library construction, which reduces handling time, reagent costs, and technical variation. However, multi-template PCRs are known to be prone to technical artifacts such as chimera formation and biased amplification [49] and the type and number of artifacts are expected to depend on the number of PCR cycles, the composition of the template pool, and the amount of template. The prime-seq protocol already minimizes the number of PCR cycles, and the composition of the template pool is given by the analysed samples and hence also not optimizable. But as we occasionally observed lower fractions of usable reads when pooling more samples, we tested whether the amount of template in the cDNA, i.e., the number of pooled samples, affects the number of usable reads.

To investigate this effect, we isolated RNA and set up reverse transcription reactions from 132 mouse brain lysate aliquots that contain ∼ 10 ng of total RNA each. We pooled 8, 32, and 92 of them for the subsequent exonuclease treatment, preamplification, and library generation, following the standard prime-seq protocol for these sample sizes. Note that this includes minimizing the number of PCR cycles, which in this case corresponds to 13, 10, and 7 cycles for the pools of 8, 32, and 92 samples, respectively.

Strikingly, the increase in input led to a decrease in the mean proportion of exonic and intronic sequences from 81.0 % to 70.1 % (Fig. 1G). In other words, using more cDNA as input in the preamplification PCR leads to fewer usable reads, despite a lower number of PCR cycles, which is considered to reduce technical artifacts in multi-template PCRs. The magnitude of this effect was surprising, also because we are not aware of any findings that report an effect of the amount of cDNA input on the quality of bulk or single-cell RNA-seq libraries. Analysing the effect in more detail, we found that the decrease in usable reads was caused almost entirely by an increase in intergenic reads and this increase was caused almost entirely by reads mapping to the opposite, i.e. antisense, strand of exons and introns, increasing from 3 % in the 80 ng condition to 11 % in the 920 ng condition (Fig. S3A). In general, antisense reads have been attributed to off-target priming, i.e., to priming of the oligodT primer and the TSO primer to other sites than the polyA tail and the 5’ end of the cDNA, respectively [50]. Off-target priming is prevalent in essentially all bulk and single-cell/single-nucleus RNA-seq libraries and a recent analysis found that on average 18 % (max. 30 %) of UMIs are antisense among the seven analyzed 10X Genomics datasets [51]. So off-target priming in general and antisense reads in particular are a common source of non-usable reads in early barcoding protocols like prime-seq. We discuss the causes and consequences of off-target priming in more detail below. Of relevance here is that there is no obvious mechanistic explanation why antisense reads are more prevalent when pooling more samples, although an increased amplification bias for shorter molecules could play a role (Fig. S3B). Independent of the mechanism, our findings indicate that pooling fewer samples is better for prime-seq and potentially also for other early-barcoding protocols. Hence, our previous recommendation of pooling up to 384 samples of 4-40 ng RNA each [16] should not be followed. Rather, we recommend that pooling should be limited to ∼ 48-96 samples with ∼ 500 ng of total RNA as input, as this seems a reasonable ad hoc compromise between the time and cost savings of pooling and the disadvantage of less usable reads.

### Fragmentation time is robust to input amounts

After its amplification, full-length cDNA needs to be fragmented to lengths compatible with short-read sequencing technologies such as Illumina. If the fragments are too long, the sequencing quality degrades [52] and clustering efficiency on the flow cell decreases [53]. Conversely, if the fragments are too short, the sequencing cycles may exceed the insert length, leading to unwanted sequencing of adapter sequences [54]. The prime-seq adapter sequences are 197 bp (including 12 bp barcode and 16 bp UMI), and we sequence the cDNA insert with only one read of 93 bp, so we aim for a total fragment size of 290-650 bp (Fig. S7). Prime-seq uses enzymatic fragmentation (NEBNext Ultra II FS) and double-sided SPRI bead purification to obtain this range of fragments. It is desirable to retain as much of the complexity present in the amplified full-length cDNA also within the fragmented cDNA, and hence to adapt fragmentation time to maximize DNA yield in the size range of 250-650 bp. We tested 5, 15, and 80 ng of full-length cDNA for fragmentation times of 2-8 minutes and determined the DNA yield in the desired size range on a Bioanalyzer (Fig. S4). We found that the 5 minutes of fragmentation time used for our original protocol works well for 5 ng and the recommended 15 ng, with a maximum yield of 66 % DNA mass at 4.5 minutes. At an input of 80 ng cDNA, fragmentation for 5 minutes yielded 60 % DNA yield and a maximum of 67 % was obtained when fragmenting for 7 minutes.

Hence, the originally recommended 5-minute fragmentation time for 15 ng cDNA works well and is reasonably robust even when 3-fold higher or lower cDNA amounts are used. Slightly longer fragmentation times are optimal for higher cDNA amounts, but as cDNA concentration is measured before fragmentation, we keep the recommendation of using 15 ng cDNA and 5-minute fragmentation also in prime-seq2.

### Improved adapter ligation and library PCR increase the number of usable reads

To generate an Illumina sequencing library from these cDNA fragments, it is necessary to add indices (i5 and i7) and the complete flow cell adapter sequences (P5 and P7) that are necessary for Illumina sequencing. As prime-seq is a 3’-tagged early barcoding method, only cDNA fragments containing the 3’ end can be utilized, as only these contain the oligodT primer with the sample-specific barcode, the UMI, and the partial P5 sequence for priming. To generate an Illumina sequencing library from these cDNA fragments, one performs end repair and dA-tailing before ligating an adapter to the 5’ end (Fig. 2A, S7). This adapter ligation generates DNA molecules that are amplified in the following “library PCR” that adds the indices (i5 and i7) and the complete flow cell adapter sequences (P5 and P7) necessary for Illumina sequencing (Fig. 2A). We suspected that the adapters and the indexing primers are suboptimal and contribute to unusable reads in the library. As interactions among adapter ligation and indexing primers are likely, we also tested the combination of these modifications. To minimize batch effects and biases, we started from one homogeneous cDNA pool and constructed three independent libraries per condition with 24 replicates each for all 12 combinations of adapters and primers, totaling 864 samples. For the indexing primers used in the original prime-seq protocol, we noticed that they might be suboptimal as their annealing sites differ considerably in length, resulting in a melting temperature (Tm) difference of approximately 6 °C. These primers were retained from prime-seq’s predecessor protocol, SCRB-seq [55], and while we are not aware of any positive consequences of this Tm difference, it is known that it can lead to inefficient and biased amplification [56]. To test whether an optimization of the library PCR could improve the number of usable reads, we tested four indexing primer combinations that differ by the presence of a read start site and hence in their length and Tm (Fig. 2A): The original protocol combines a long P5 primer that contains the read start site of Illumina’s Read 1 sequencing primer (”P5+RS”) and a short P7 primer that does not contain the read start site of Illumina’s Read 2 sequencing primer (”P7-RS”). We designed the two additional versions, P5-RS and P7+RS, resulting in the four possible primer combinations. We adjusted the annealing and elongation temperature from 65 °C to 72 °C for the P5+RS and P7+RS primer combination, and generated libraries from the same mouse brain cDNA.

**Figure 2.**
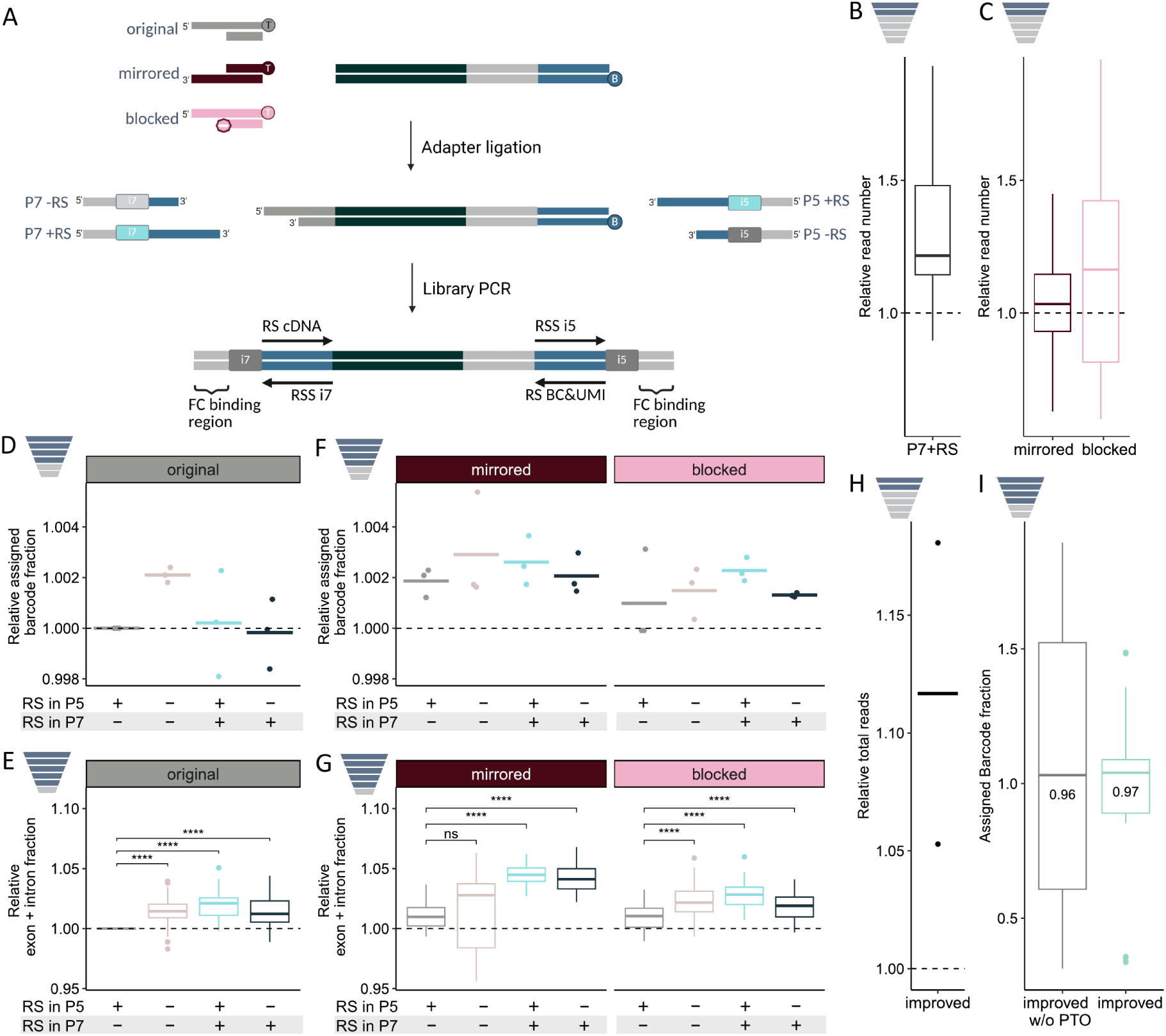
Possible improvements during adapter ligation and library PCR. **A**, Principle of adapter ligation and library PCR. Adapters are ligated to the cDNA fragments after fragmentation using a 3’ A-overhang. Note that the biotinylated SINGV6 primer prevents ligation on the P5 site (B - Biotin). Alternative mirrored and blocked ligation adapters were designed to prevent the formation of P7-P7 fragments in the following library PCR that amplifies fragments and adds indices (i5 and i7) as well as the complete flow cell adapter sequences (FC binding regions P5 and P7). In the original protocol, a P5 primer with and a P7 primer without a read start (P5+RS & P7-RS) were used, and we tested all four primer combinations with and without a read start for all three adapters with 24 samples per condition and three independent library preparations each (n= 864). **B**, Median ratio of read numbers after index assignment with the new P7+RS relative to the original P7-RS. (n=18 pairs). **C**, Median ratio of read numbers after index assignment with each new ligation adapter relative to the original ligation adapter. (n=24 pairs each). **D**, Mean fraction of assigned barcodes per combination of index primers with the original ligation adapter (n=3 independent library replicates per condition) **E**, Mean fraction of mapped exonic and intronic reads per combination of index primers with the original ligation adapter (n=72 samples per condition) **F**, Mean fraction of assigned barcodes per combination of index primers with the new ligation adapters (n=3 independent library replicates per condition) **G**, Mean fraction of mapped exonic and intronic reads per combination of index primers with the new ligation adapters (n=72 samples per condition) **H**, Total reads relative to the improved protocol without PTO bonds (n=2 independent library replicates per condition. Bar shows the mean; error bars show maximal/minimal values.). **I**, Fraction of assigned barcodes with and without PTO bonds (n=42; Two independent library replicates of 10/11 samples per condition.). P-values were calculated in paired two-sided t-tests (ns: p *>* 0.05; *: p ≤ 0.05; **: p ≤ 0.01; ***: p ≤ 0.001; ****: p ≤ 0.0001).

We find that libraries generated with P7+RS instead of the original P7-RS have 37 % more reads with a correct i5 and i7 index (index assignment step), and libraries generated with P5-RS instead of P5+RS have 43.2 % fewer (Fig. 2B, S5). Hence, the longer primers generate more sequenceable library molecules, confirming that our previous conditions were suboptimal. Importantly, these additional reads generated by P7+RS and P5+RS library PCRs had the same fraction of reads with assigned barcodes (Fig. 2D) and even a 2 % higher fraction of reads in exons and introns (Fig. 2E), indicating that not only more, but also more usable reads are produced when using P7+RS and P5+RS indexing primers. The ligation adapter used in the original prime-seq protocol has a 15-base 5’-overhang, so that the P7 index primer can anneal only after the P5 index primer has elongated a fragment (Fig. 2A). This setup is intended to prevent the amplification of P7-P7 fragments that are generated from internal cDNA fragments that have ligation adapters on both ends. However, this design overlooked that the polymerase can synthesize the complementary overhang of the ligation adapter, enabling the priming of P7 index primers without the prior elongation of the P5 index primer.

To address this issue, we developed two variants of ligation adapters (Fig. 2A). The first variant is a mirrored adapter, which has a 3’-overhang instead of a 5’-overhang. This variant leads to complete amplicons only when molecules are initially primed and elongated from the P7 site. Initial priming and elongation from the P5 site, as well as P7-P7 fragments, generate ”dead-end” amplicons lacking an overhang on the opposite side for P5 binding. The second variant is a blocked adapter, which includes a dideoxycytidine at its 3’-end. This prevents the synthesis of the complementary overhang and hence should lead to the preferential amplification of fragments with a P5 and a P7 end.

The mirrored and the blocked adapters increased the number of reads after index assignment by 5 % and 16 %, respectively (Fig. 2C). As for the increased number of reads produced by P7+RS compared to P7-RS, this improvement is likely due to a reduction in irregular fragments, which are quantified but fail to produce valid reads. Importantly, these additional reads with ∼2 % also had a slightly higher fraction of reads with barcodes (Fig. 2F) and an up to 5 % increase in exonic and intronic reads (Fig. 2G), indicating that not only more, but also more usable sequences are produced.

In summary, both new ligation adapters perform better than the original ligation adapter regarding the number of sequencable reads, fraction of assigned barcodes, and fraction of mapped exons and introns. Both perform best when combined with the P7+RS & P5+RS index primers. As the blocked adapter performed slightly better with respect to the number of sequenceable reads and because it prevents, by design, more mispriming than the mirrored adapter, we integrated the blocked adapter and the P7+RS & P5+RS index primers into the prime-seq2 protocol.

### PTO bonds prevent primer editing

Proofreading polymerases are commonly used in RNA-seq protocols due to their 3’-5’ exonuclease activity, which reduces PCR errors [57, 58, 59]. However, when mismatches occur between the primer and template, these polymerases can modify primer sequences, leading to unspecific amplification [60, 61]. Previous studies have shown that incorporating phosphorothioate (PTO) bonds into the primers’ 3’ ends can block this primer editing [62, 60]. We incorporated PTO bonds into the last two 3’ bonds of all PCR primers (SINGV6, P5+RS, P7+RS) and tested their effects using the lysates from mouse striatal punch biopsies.

We detected a mean increase in total reads of 11.7 % when using primers with PTO bonds (Fig. 2H). This could be attributed to a decrease in irregular fragments resulting from mispriming, which distort quantification but do not produce valid reads as described above. We also noted a 1.4 % higher fraction of mean assigned barcodes with significantly less variation among samples (Levene’s Test, p*<*0.05; Fig. 2I).

Hence, the incorporation of PTO bonds improved the number of usable reads, and we integrated primers with PTO bonds in the prime-seq2 protocol.

### Validation of the prime-seq2 protocol

Finally, we compared the combination of improvements in the protocol with the original prime-seq protocol using lysates from mouse striatal punch biopsies, generating four libraries of 32 samples per protocol. The improved protocol (prime-seq2) includes the optimized DNA digest, the 3’ blocked ligation adapter, a library PCR with P5+RS and P7+RS primers, and their higher annealing temperature and PTO bonds in SINGV6, P5+RS, and P7+RS primers. Note that for both protocols, 32 samples are pooled, and potential additional improvements from pooling fewer samples (Fig. 1G) are not included in this comparison.

For both protocols, we visualize the fraction of reads remaining after each step of the read funnel, arbitrarily setting the average number of total reads obtained from the prime-seq2 libraries as 100 % (Fig. 3A). What matters most is that the number of usable reads at the end of the funnel, i.e., the fraction of UMIs, increases from 28 % to 45 %, i.e., by 60 % (Fig. 3A, E). Hence, the prime-seq2 protocol represents a considerable improvement in the number of usable reads. Going through each step of the funnel for the total read fraction (Fig. 3A) and for the relative change (Fig. 3B) helps to explain how this improvement occurs.

**Figure 3.**
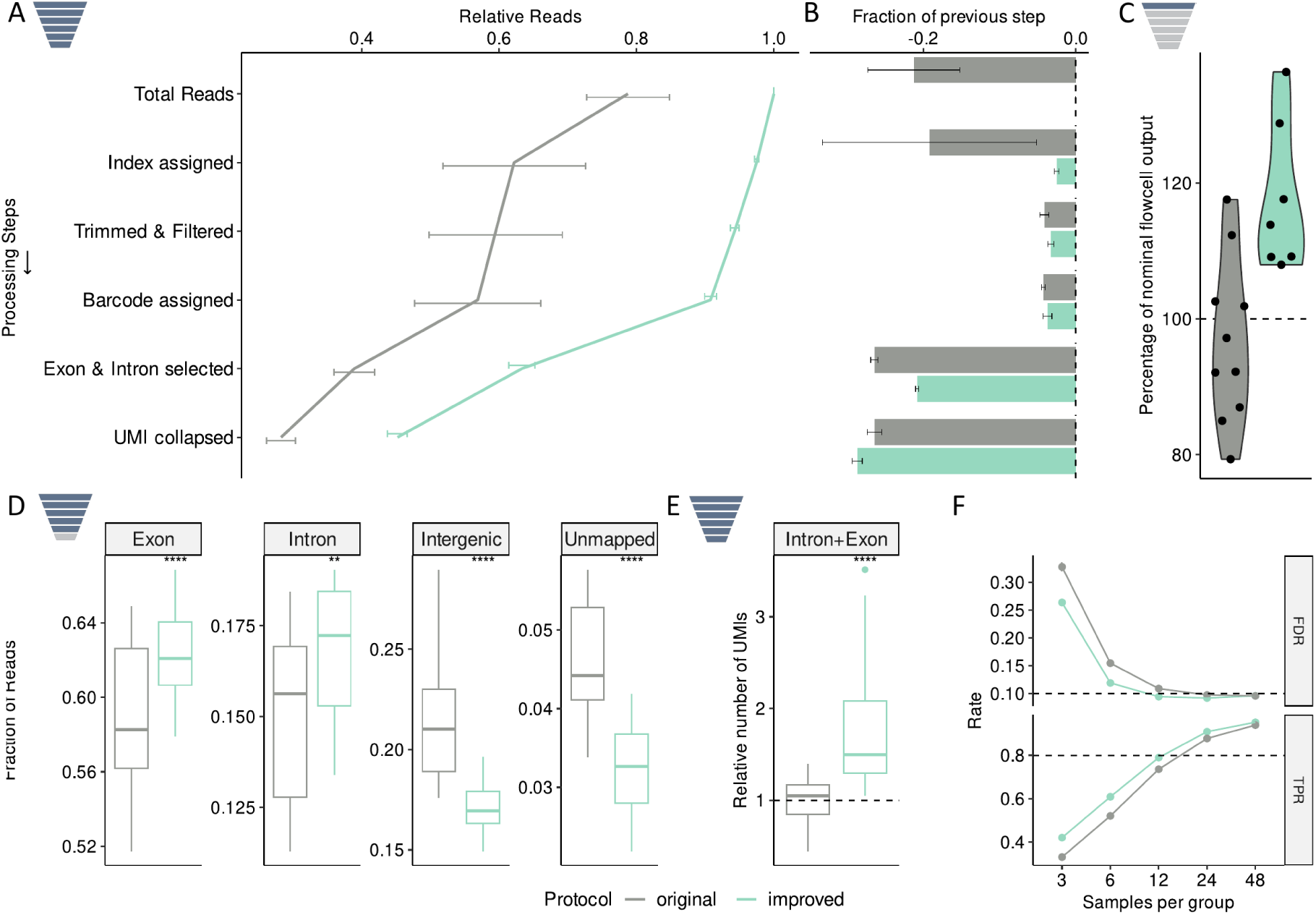
Overall comparison of the improved protocol. **A**, Funnel plot showing the read number at each step relative to the total reads with the improved protocol. The first four steps use one data point per library, while the last two show one data point per sample, increasing the variation. Lines show means; error bars show SE. (n=64; Four libraries of 32 samples (8 barcodes) per condition.) **B**, Change in total reads per step relative to the previous step. Error bars show SE. **C**, Total reads per flow cell relative to the nominal flow cell output according to the manufacturer. (n=17) **D**, Fraction of barcode-assigned reads mapped to exons, introns, or intergenic regions and unmapped reads. (n=64; Four libraries of 32 samples (eight barcodes) each per condition.) **E**, Number of UMIs relative to the original protocol. (n=64; Four libraries of 32 samples (eight barcodes) each per condition.) **F**, Power analysis of both protocols at equal flow cell share across replicate numbers. Points and lines show means; error bars show SE. P-values were calculated in unpaired two-sided t-tests (ns: p *>* 0.05; *: p ≤ 0.05; **: p ≤ 0.01; ***: p ≤ 0.001; ****: p ≤ 0.0001).

The new protocol prime-seq2 increases the total reads by a mean of 27 %, i.e., for a given amount of library molecules, 27 % more reads are obtained from a flow cell. As explained above, we think that this is caused by fewer molecules that get quantified but fail to cluster or - more importantly - cluster, but fail to produce an R1 sequence read and hence do not get detected by the sequencer. Of note, we observed this effect not only in the direct comparison of the two protocol versions done here, but also across 17 unrelated projects from a variety of species and cell types that we sequenced on Illumina’s NextSeq1000/2000 using P2, P3, and P4 flow cells (Tab. S1). The 10 and 7 libraries prepared with the old and the new protocol had 97 % and 118 % of the expected flow cell output, respectively (MWU test, p*<*0.05; Fig. 3C). In the next processing step - index assignment - 19 % of the reads get filtered out in the original protocol, but only 2.5 % in prime-seq2. This indicates that more molecules contain a proper i5 and i7 index in prime-seq2. Both protocols then filter a similarly small percentage of reads during trimming & filtering (3-4 %) and barcode assignment (4 %). This is partly expected, as neither the RT-primer nor the fragmentation and size-selection were changed in prime-seq2, and partly indicates that the PTO bonds do not improve the assigned barcode fraction decisively. The generally small percentage of filtered reads shows that these two steps are not a major bottleneck for the number of usable reads in prime-seq. A considerable fraction of barcode-assigned reads, 26 % and 21 % for the old and the new version, respectively, is filtered because they do not map to an annotated exon or intron. 4 % and 3 % cannot be mapped at all and 21 % and 17 % map to intergenic regions (Fig. 3D). Hence, prime-seq2 does not only produce 60 % more barcode assigned reads (Fig. 3A), but among them is a significantly higher fraction of exonic and intronic reads. This is probably caused largely by the improved DNA digest (Fig. 1D) and the new index primers (Fig. 2E). Why, despite a presumably complete DNA digest, the fraction of intergenic reads is still at 17 % is interesting and discussed in more detail below. In the final UMI collapsing processing step, 26 % and 29 % of the exonic and intronic reads are filtered out. Hence, there is a slightly higher fraction of duplicated reads in the prime-seq2 protocol for unclear reasons. Importantly and as mentioned in the outset, the final number of usable reads, i.e. the total number of exonic and intronic UMIs increases significantly from 28 % to 45 %, i.e. by 60 % (Fig. 3A, E).

To quantify how this improvement translates into performance, we used power simulations as implemented in powsimR [30]. PowsimR estimates the mean-variance distribution of the data, in our case, of the two protocol versions, and then simulates differently expressed genes between two conditions to obtain a ground truth against which simulations for a certain number of replicates per condition can be compared. We estimated the mean variance distributions using equal flow cell shares and hence took the increased number of total reads of prime-seq2 into account. We simulated that 10 % of genes were differentially expressed, with log-fold changes following a gamma distribution, and varied the number of replicates per condition from three to 48. The prime-seq2 protocol consistently had a higher true positive rate (TPR) in detecting differentially expressed genes across all numbers of replicates. prime-seq2 achieves an 80 % TPR with only 13 samples per group, whereas the original prime-seq protocol requires 17 samples to reach the same threshold. Importantly, the false discovery rate (FDR) was not compromised and was actually lower than that of the original protocol for low replicate numbers and similar at higher replicate numbers (Fig. 3F, S6). Therefore, the protocol improvements increase the power to detect differentially expressed genes without additional sequencing costs.

## Discussion

Bulk RNA-seq is a powerful method for many biological questions, and its cost-efficiency is a crucial factor, limiting biological insights at a given budget. Early barcoding protocols have reduced library costs per sample by 10-50-fold by minimizing reagent costs and increasing throughput [16]. What has been largely neglected in protocol developments is understanding and improving the number of usable reads that are obtained from RNA-seq libraries. Here, we systematically optimized the 3’ tagged early barcoding protocol prime-seq. This results in the improved protocol prime-seq2 that generates 60 % more usable reads and presents one of the most widely used and cost-efficient home-brew bulk RNA-seq protocols available. Further, we think our optimization approach reveals insights and approaches how to optimize other bulk, single-cell, and single-nucleus RNA-seq protocols. Finally, our results provide insights into the types of molecules that get amplified in such protocols and how this affects the interpretation of RNA-seq profiles. We discuss these issues in turn.

### Powerful and cost-efficient RNA-seq with prime-seq2

While using the original prime-seq protocol [16] in various projects, we noticed that some samples generated fewer usable reads than expected, especially after switching to sequencing on Illumina’s patterned flow cells. We identified three main issues: (i) lower-than-expected total sequencing reads per flow cell, (ii) loss of reads due to missing index assignments, and (iii) a high proportion of intergenic reads. By systematically testing adjustments of different protocol steps as presented above, the resulting prime-seq2 protocol led to 27 % more reads per flow cell, 21 % more index assignments and 7.6 % more exonic and intronic reads (Fig. 3A-C). While it is not possible to unambiguously attribute each improvement to specific adjustments, we propose the following plausible explanations:

(i): Lower-than-expected total sequencing reads can be obtained when library molecules are quantified, but fail to amplify on the flow cell, and when library molecules get amplified, but lack a functional Read 1 sequencing primer binding site (Fig. 2A, S7). Our improvement of 27 % (Fig. 3A-C) is likely caused by blocked ligation adapters, longer P7 primers and the primers with PTO bonds as these modifications are expected to increase the specificity of the library PCR and hence produce more sequenceable molecules. This is supported by our findings that these modifications lead to 16 % (Fig. 2C), 37 % (Fig. 2B), and 12 % (Fig. 2H) more index-assigned reads, respectively. (Note that these numbers reflect both the increased total reads and the improved index assignment. Disentangling these two factors would require separate flow cells, which was only done during validation of the full protocol (Fig. 3A-C)). (ii): Clusters that get filtered out at the index assignment steps have either no proper i5 or no proper i7 index. Similar to library molecules that lack a functional Read 1 primer binding site, this can be caused by molecules lacking a functional Read 2 (i7) sequencing primer binding site or lacking a functional Read 3 sequencing primer binding site (i5). As above, our improvement of 21 % (Fig. 3A-B) is likely caused by our blocked ligation adapters (Fig. 2C), longer P7 primers (Fig. 2B), and the incorporation of PTO bonds (Fig. 2H), as these should increase the specificity of the library PCR and hence produce more sequenceable molecules. Importantly, this would help little if the additional reads did not contain usable reads mapping to exons and introns. Fortunately, the improved library PCR generates even 3 % more of those reads (Fig. 2G) and hence really does increase the number of usable reads. (iii) Intergenic reads, i.e., reads mapping to the genome, but not to the sense strand of annotated exons and introns, were reduced from 21 % to 17 % in the improved protocol (Fig. 3D). Intergenic reads were reduced by optimizing DNA digest conditions, which seems especially relevant if degraded genomic DNA is present (see Fig. 1C-D, 2). As no genomic DNA is detectable after the optimized digest (Fig. 1C), essentially all intergenic reads in the improved protocol must be derived from RNA. Remarkably, the fraction of intergenic reads can vary considerably and is influenced by the conditions of reverse transcription (Fig. 1E), the amount of samples pooled for cDNA amplification (Fig. 1G), and by the library PCR conditions (Fig. 2G). The causes and consequences of intergenic reads are discussed in more detail below. Here, it is worth mentioning that the biggest reduction of over 10 % of intergenic reads is achieved when pooling 8 lysates à 10 ng total RNA instead of 92 (Fig. 1G). Note that this effect is masked in our final protocol comparison, as we pooled 32 samples both for the original as well as the prime-seq2 protocol. We currently do not know how this pooling effect is caused, but the increased amount of antisense reads and the reduced representation of long transcripts (Fig. S3) indicate that pooling more cDNA could increase the amplification bias towards shorter molecules in the preamplification and/or the library PCR. This likely plays a role also in other early barcoding bulk and scRNA-seq protocols and hence might be a relevant parameter that can improve early barcoding protocols in general. For the current version of prime-seq2, we recommend limiting the libraries to pools of ∼500 ng of total RNA, which is a reasonable compromise for most of our projects with respect to intergenic reads, reagent costs, and hands-on time. In summary, prime-seq2 delivers on average 60 % more UMIs than prime-seq and hence reduces sequencing costs by an average of 38 %. For example, this allows for the generation of bulk RNA-seq libraries of 96 samples at 2.5€ per sample, including RNA isolation (Tab. S2) and sequencing data for another 20-40€ per sample (assuming 5-10 million usable reads per sample and 2€ per million paired-end reads). To our knowledge, this makes it the most cost-efficient bulk RNA-seq protocol [16], although other early-barcoding bulk RNA-seq protocols are in the same range, and proper benchmarking would be required to compare cost-efficiency quantitatively. Prime-seq is well used by us and others [63, 64, 65, 66, 67, 68, 69] and due to its low set-up costs, which essentially comprises the 96 custom oligos for ∼1250 € (Tab. S2), it is an attractive alternative to commercial kits for core facilities and labs that frequently perform bulk RNA-seq. The detailed step-by-step protocol of prime-seq2 is accessible on protocols.io: doi.org/10.17504/protocols.io.14egn97kpl5d/v1.

### Monitoring and optimizing RNA-seq protocols using the funnel

Our strategy to monitor and optimize prime-seq could be useful also for other RNA-seq protocols, as the number of usable reads varies widely within and among other bulk and single-cell RNA-seq protocols [25, 70] and very similar principles like poly(A) priming, early barcoding, and cDNA and library amplification apply to most of them. We think that just monitoring the different filter steps from total reads to usable reads in more detail, i.e., by applying the funnel strategy, could be a useful QC metric that can identify issues for particular samples, protocols, and/or sequencing platforms. For example, lower rates of index assignment can pinpoint non-sequenceable library molecules, or a high fraction of intergenic reads could pinpoint suboptimal library amplification, as discussed above for prime-seq. If appropriate, identified bottlenecks could then be addressed by adjusting individual protocol steps. When optimizing such steps, it is important to also track other performance parameters such as complexity or technical noise as was e.g. relevant when we tried optimizing RT-conditions or for prime-seq (Fig. 1F). It can also be important to consider non-obvious interactions of sample types and protocol steps as we found for the DNA digest that was apparently working well for samples with high molecular weight genomic DNA contamination [16], but turned out to be optimizable for samples with degraded genomic DNA modifications (Fig. 1D). Another illustration of the usefulness of the funnel is that for prime-seq2, still about 29 % of the reads are lost in the UMI-collapsing step (Fig. 3A). This indicates that measures to increase the complexity of the library, such as increasing the input further, optimizing cycle number, and amplification conditions, might be a bottleneck that is worth addressing next.

In summary, we think that the funnel can be a useful QC metric to monitor and optimize bulk and scRNA-seq protocols.

### Priming, mispriming, and library complexity

Our optimizations also shed light on another aspect of polyA-primed RNA-seq protocols that we think is worth discussing, which is the relation between the specificity of the oligo-dT priming and the complexity of the resulting libraries. The idealized view is that oligodT priming on the polyA tail of an mRNA leads to reverse transcription of the (entire) mRNA and is followed by template switching and second-strand synthesis. The following cDNA amplification and library PCR leads to library molecules in which - for 3’ tagged protocols -the 3’-end of the cDNA is read, starting ∼200bp downstream of the polyadenylation site (Fig. 1B and 7). Due to only one priming site - the polyA tail - the number of reads is independent of the length of the mRNA. However, stretches of adenine also occur within transcripts (”internal polyA”), which inevitably leads to internal oligo-dT priming, for which already six or seven consecutive adenines seem sufficient [71]. This off-target priming is prevalent in bulk and single-cell RNA-seq data and ranges from *<*5 % - 65 % for exonic reads and *>*95 % for intronic reads, dependent on the protocol and the sample (Table S1 in [71]). As intronic reads are derived largely from nascent, i.e., (partially) unspliced transcripts, and those are confined to the nucleus, the use of intronic reads is especially relevant for single-nucleus RNA-seq [72, 27] ). But also for single-cell data, intronic reads can account for over 30 % or over 50 % of all reads from mouse embryonic brain samples or human PBMCs, respectively [50]. This creates a dependency between the number of reads and the occurrence of internal poly(A) stretches that in turn leads to the observed gene length bias, i.e., a correlation of number of reads and transcript length [73, 74, 71]. This reduces the correlation of read counts and the number of RNA molecules in the samples, i.e., reduces the accuracy of the RNA quantification. It has been proposed to filter or correct these off-target reads [71], but this does not work well for real data [74]. However, most bulk and single-cell analyses do not profit from a slightly higher accuracy, as they focus on comparing relative RNA levels among samples or cells to identify differentially expressed genes or distinguish different cell types. For those purposes, the complexity of a library is much more relevant, i.e., how many genes are detected at a given sequencing depth or - more precisely - for how many genes one has sufficient reads, i.e. statistical power, to detect differential expression. As it turns out that thousands of genes are largely or even exclusively detected by intronic reads [75], off-target reads are now considered more as a feature than as a bug, exemplified by intronic reads being now the default for single-cell and single-nucleus data in the 10x CellRanger software [76]. In the context of this study, this matters, as we not only assumed that intronic reads are usable, but we also observe a relationship of priming specificity and library complexity: When increasing the temperature for the reverse transcription reaction, we do increase the number of usable, i.e. exonic and intronic reads by 6 % (Fig. 1E), but we also decrease the complexity by detecting only 2/3 of the genes (Fig. 1F). As alluded to above, this can be explained if the higher temperature reduces off-target priming for many genes that are not well detected by on-target oligo-dT priming. Also, other protocols probably differ considerably in off-target priming rates as seen by the differences in library complexity and fraction of intronic reads among commercial scRNA-seq protocols [70]. For example, the 5’ expression kit from 10x has three times fewer intronic reads than the 3’ expression kit [70, 50]. This is likely caused by the lower off-target priming rate of the TSO compared to the oligodT [50] and this likely leads to the lower number of detected genes per cell (∼2000 and ∼3000 in 5’ and 3’ PBMC data, respectively [70]), as a similar reduction in gene number is seen when counting solely exons in 3’ expression data [76]. While the balance between off- and on-target priming seems important for the performance of RNA-seq protocols, it is currently unclear where an optimal balance might lie, how it could best be measured, and whether it should and could be optimized. In any case, it is likely that it is a major factor contributing to differences in gene expression levels among protocols [22], among RNA isolation methods [16] , or usable reads as discussed below. In summary, we find for prime-seq that more stringent reverse transcription leads to slightly more usable reads, but less complex libraries. This stresses the general importance of optimizing not one parameter in isolation and likely reflects the balance between off- and on-target priming, which is an important but largely unexplored parameter for RNA-seq libraries.

### Usable and unusable reads in RNA-seq

Barcode assigned reads (”reads” in the following) are mapped to the genome and are used further if they map to the sense strand of an annotated exon or intron. For the most recent mouse gene annotation GENCODE34 [77] that is used here, these are 21,414 annotated protein coding genes, 15,131 long non-coding RNA genes, 6,105 small non-coding RNA genes or 13,754 pseudogenes that amount to 19.5 % of the mouse genome. A considerable fraction of reads - 17 % for the prime-seq2 validation data (Fig. 3D) - map to intergenic regions and their origin and information content seem worth discussing. Several possible sources for these intergenic reads can be distinguished: 1) Genomic DNA: This is a potential source of intergenic reads as the reverse transcriptase can use DNA as template. However, as discussed above, this source is probably negligible for prime-seq2 due to its optimized DNA digest. 2) Mismapping: A substantial fraction of reads in RNA-seq - ∼10 % of reads for our data - map equally well to 2-50 different regions of the genome (*>*50 are filtered out). There are different possibilities to handle such multimapped reads [78] and STAR randomly assigns them to one region. Among the intergenic reads, 2.9 %pt. have an equally good match in an exonic or intronic region (data not shown). Assuming that such reads actually do originate from an annotated gene - as currently assumed, e.g., by STARsolo - this would reduce the fraction of intergenic reads to 14.1 %. 3) Antisense: For the 3’tagged libraries of 10x, it has been convincingly argued that antisense reads arise when the second cDNA strand is not generated by the TSO, but instead by the oligodT primer hybridizing to internal polyA stretches on the first strand cDNA [50]. This generates one sense and one antisense molecule in the library, indeed leading to a correlation of sense and antisense reads that is even higher than the exon-intron correlation [50]. For 10x scRNA-seq data, on average 14 % of all UMIs mapping to exons/introns are antisense [50]. For prime-seq2, we find comparable numbers, with ∼5 % of all reads being antisense (Fig. S3A), which corresponds to ∼9 % of all UMIs mapping to exons/introns. To our knowledge, it has not been investigated whether using antisense reads would increase or decrease the power to detect differentially expressed genes in bulk or scRNA-seq data, but as introns do increase this power, the even more correlated antisense reads are likely to do so as well. However, the current practice, as evident by the default settings of CellRanger and STARsolo, is not to use them. Of note, RNA-seq protocols based on random priming are not strand-aware and hence necessarily use sense and antisense reads to quantify expression levels. 4) Non-annotated read-through isoforms: It has been shown that thousands of genes that are detected by in situ hybridization in a mouse brain region have reads from a 10x experiment of the same region mapping within 10 kb downstream of their annotated 3’ end [75].

These reads make up 25 % of all intergenic reads in this dataset and ∼2000 genes are only detected when using these reads [75]. A good proportion of these reads might be derived from short-lived read-through transcripts that arise especially during stress [79], but read-through transcripts are also found for 1/3 of all protein-coding genes in ”normal” human tissues [80, 81, 82]. Either way, they can be regarded as non-annotated isoforms of the upstream genes and, similar to using introns, increase the effective complexity of the library by allowing more genes to be tested for differential expression. Also in our data, we find ∼25 % of all intergenic reads to map in sense orientation to regions 10 kb downstream of annotated genes (data not shown), suggesting that a good fraction of intergenic reads might be derived from non-annotated read-through transcripts. While it has been suggested to incorporate such reads for quantification [75] and recent ENCODE versions have taken this up and integrated options for extending human and mouse gene annotation [83], it is not the current practice to use those reads. 6) Pervasive transcription: After microarrays and Next-Generation Sequencing had made deep transcriptome profiling possible, it became clear that a large fraction of the genome is transcribed to some extent [84]. This includes the above read-through transcripts, transcripts originating from enhancers (eRNA) or promotors (PROMPTS), and gives rise to the 36,172 currently annotated long non-coding RNAs [83] and many more non-annotated ones [85]. The overly enthusiastic functional interpretations of these transcripts have been rightly criticized [86] and it is not clear how to interpret annotated and non-annotated lincRNAs in quantitative RNA-seq experiments. Many probably do reflect the transcriptional activity of the region in some way, and interestingly, it has been suggested to exploit these reads in scRNA-seq data to infer chromatin states [51].

In summary, the 17 % intergenic reads consist of 2.3 %pt. mismapped reads, ∼5 %pt. antisense reads, and 4.3 %pt. putative read-throughs. Most of these reads are probably directly related to the canonical transcription of (protein-coding) genes and hence probably improve their detection and quantification, similar to introns. However, in contrast to introns, there is no consensus when and how often these reads might originate from different processes, like stress-induced read through or genuine antisense transcription that might regulate sense transcription [87]. It is probably due to this uncertainty that they are not used in current practice. However, given that they do help to detect and quantify genes known to be expressed [75], further work is warranted to explore whether this practice might be worth changing.

## Conclusion

In this study, we have optimized the number of usable reads by systematically testing different steps in a bulk RNA-seq protocol. This has resulted in prime-seq2, one of the most cost-efficient home-brew bulk RNA-seq protocols available. Our study suggests that monitoring the filtering of usable reads can serve as a valuable quality control for many bulk and single-cell RNA-seq protocols. It also highlights the complexity of the conditions and interactions that shape RNA-seq library composition, with implications for how these data should be interpreted.

## Data Availability

The data discussed in this publication have been deposited in NCBI’s Gene Expression Omnibus [88] and are accessible through GEO Series accession number GSE286365 (ncbi.nlm.nih.gov/geo/query/acc.cgi?acc=GSE286365). The remaining data and code for data analysis and visualization can be accessed at github.com/Hellmann-Lab/prime-seq2 and as a stable version at 10.5281/zenodo.16902725

## Supplementary Data

Supplementary Data are available at NAR Online.

## Author Contributions

FP, EB, WE and DR conceived the study. FP, EB and DR conducted the experiments. FP and EB analyzed the data. FP, EB, WE and DR wrote the manuscript.

## Acknowledgments

We thank Ines Bliesener for lab support and the Enard/Hellmann group for helpful discussions. We thank Dr. Stefan Krebs and the staff of LAFUGA for sequencing services. Some illustrations in the graphical abstract, Fig. 1-3, and Fig. S7 were created with BioRender.com.

