## Supplemental Figures for "Improving RNA-seq protocols"

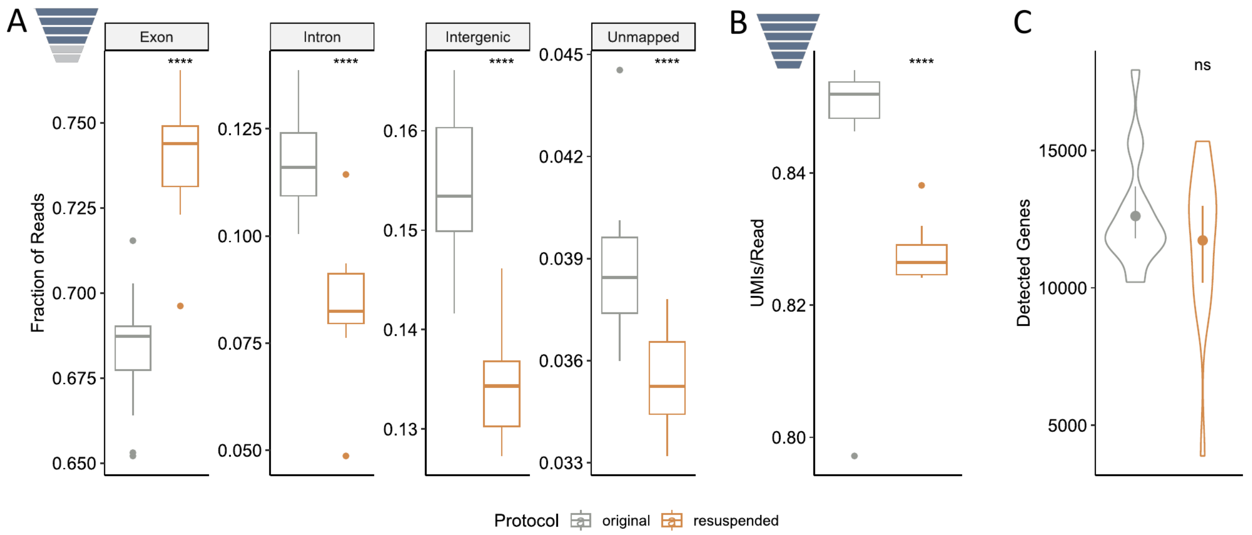


**Supplemental Figure 1. Testing of additional resuspension steps. A**, Fraction of barcode-assigned reads mapped to exons, introns or intergenic regions and unmapped reads. (n=32; 16 samples per condition in one library.) **B**, UMI count per read showing the loss of reads during UMI collapsing. (n=32; 16 samples per condition in one library.) **C**, Complexity *i.e.* the number of detected genes after downsampling to 1.25 million reads per sample, showing a slight but non-significant reduction. (n=32; 16 samples per condition in one library.)
P-values were calculated in unpaired two-sided t-tests (ns: $p > 0.05$; *: $p \leq 0.05$; **: $p \leq 0.01$; ***: $p \leq 0.001$; ****: $p \leq 0.0001$).

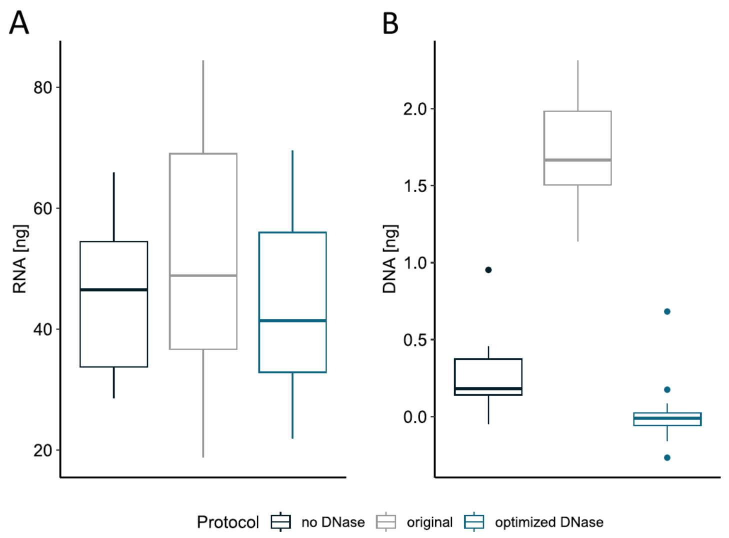


**Supplemental Figure 2. More extensive plots on DNase tests.** DNA digest was compared using the original protocol and an optimized DNA digest. **A**, The plot shows RNA amount per condition after Proteinase K and DNase I treatment. (n=96; Two independent replicates of 16 samples each per condition.) **B**, DNA amount between protocols including the negative control without DNase, showing the unintended DNA removal by SPRI bead clean-up. (n=96; Two independent replicates of 16 samples each per condition.)

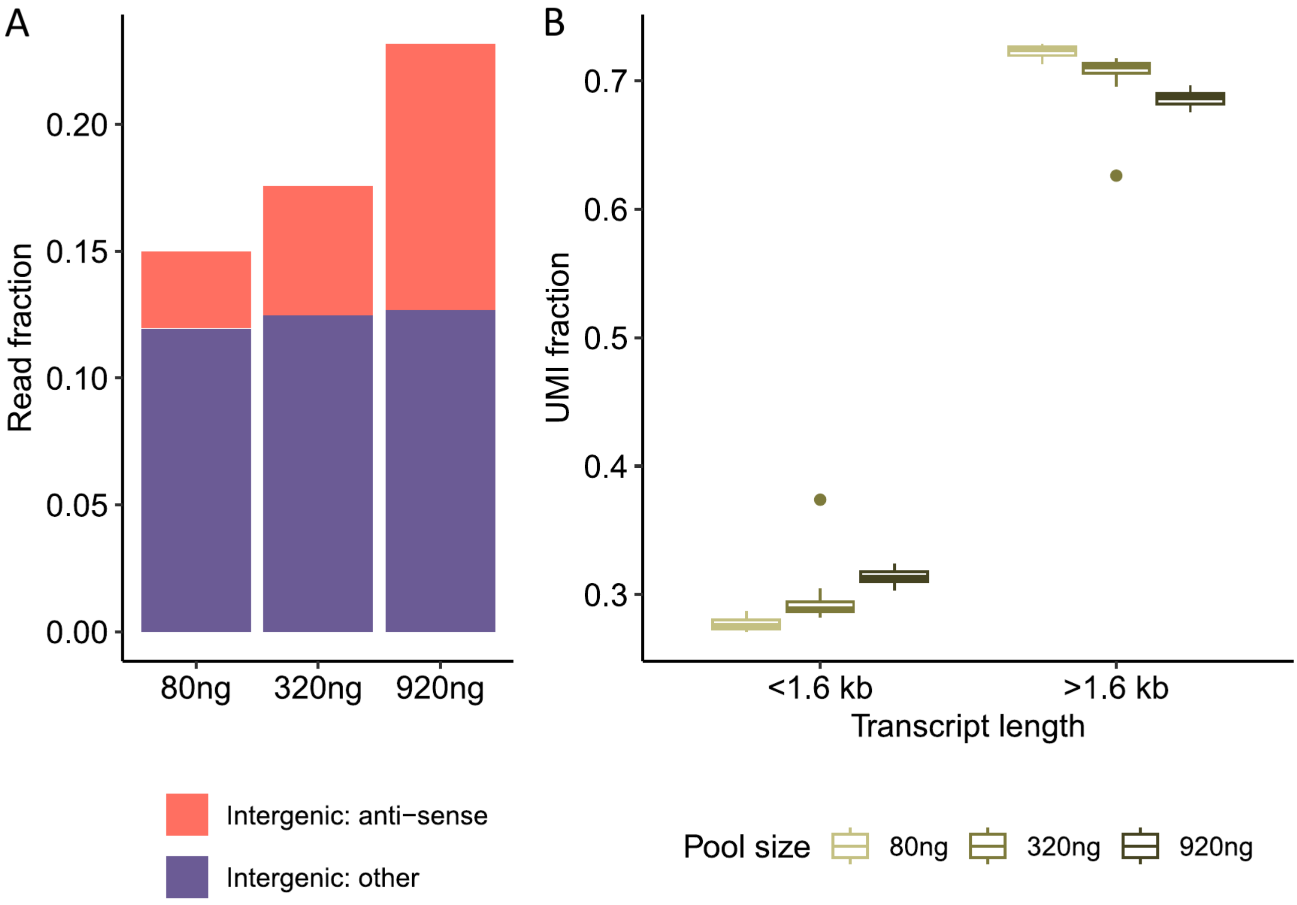


**Supplemental Figure 3. cDNA pool size increases PCR length bias and anti-sense reads.** The effect of increasing cDNA pool size on PCR amplification bias and generation of anti-sense reads was evaluated. **A**, UMI counts were categorized based on transcript length (¡1.6 kb and ¿1.6 kb). Increasing pool size elevates the fraction of UMIs from shorter transcripts (¡1.6 kb) and decreases the fraction from longer transcripts (¿1.6 kb). (n=8 for 80 ng and n=16 for 320 ng and 920 ng) **B**, Intergenic reads are subdivided into those mapping anti-sense to annotated genes and those mapping elsewhere (other). Anti-sense intergenic read fractions increase with greater pool size, whereas other intergenic reads remain unchanged.

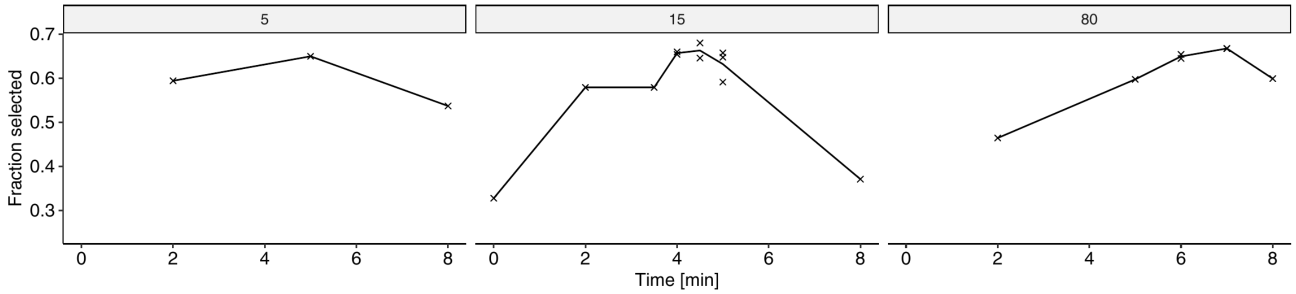


**Supplemental Figure 4. Comparison of fragmentation times.** Fraction of DNA amount in size selection range (250-650 nt) of total DNA amount (100-8000 nt). (n=21)

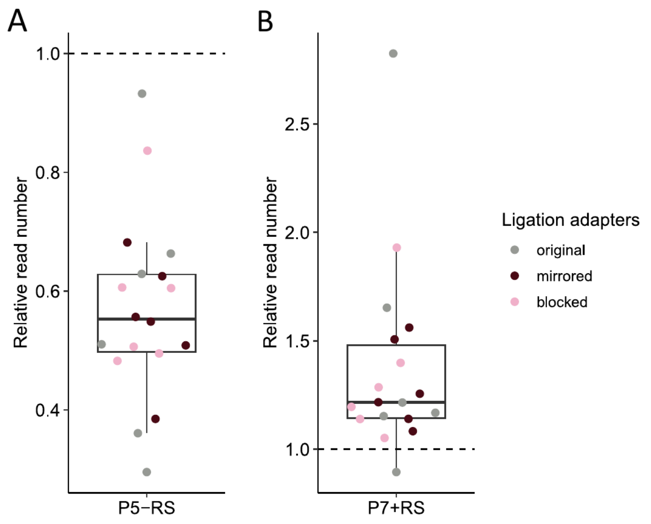


**Supplemental Figure 5. Effects of index primer length on read numbers.** DNA digest was compared using the original protocol and an optimized DNA digest. **A**, Number of reads with the new P5-RS compared to the original P5+RS. Coloring by ligation adapter shows consistent effects across modifications. (n=18; Three independent library replicates of six samples per condition.) **B**, Number of reads with the new P7+RS compared to the original P7-RS. Coloring by ligation adapter shows consistent effects across modifications. (n=18; Three independent library replicates of six samples per condition.)

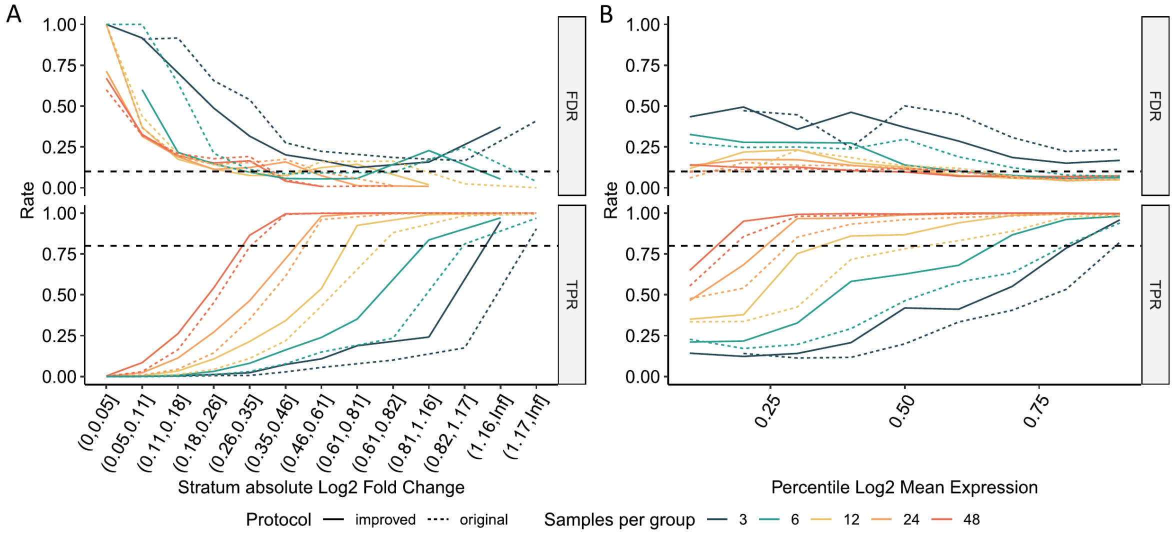


**Supplemental Figure 6. Power analysis stratified by log fold change or mean expression.** Power analysis of the original and improved prime-seq protocol at equal flow cell share across replicate numbers **A**, stratified by absolute log fold change and **B**, stratified by log mean expression.

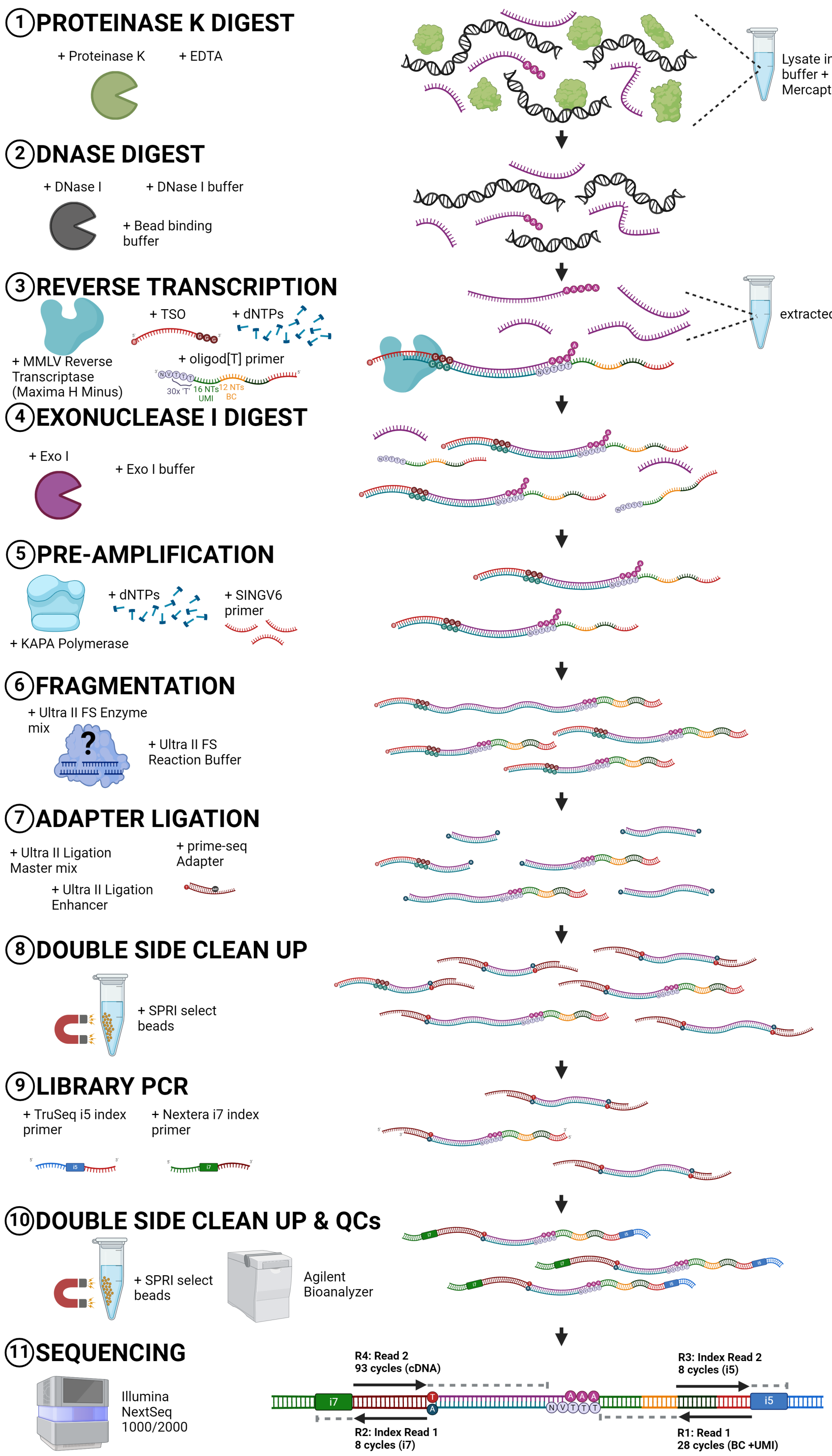


**Supplemental Figure 7. prime-seq workflow.**
